# Single-Nuclei Transcriptomic Characterization of *APOE4*-Associated Alzheimer’s Disease

**DOI:** 10.64898/2026.04.03.715591

**Authors:** Kaitlin Murtha, Anjalika Chongtham, Won-Min Song, Marjan Ilkov, Minghui Wang, Catherine Qing Chen, Claudia De Sanctis, Risit Datta, Dushyant Purohit, Edward B. Lee, Bin Zhang, Ana C. Pereira

## Abstract

Apolipoprotein E (*APOE*) genotype contributes significantly to Alzheimer’s disease (AD) risk and pathogenesis. Cell-type specific effects of *APOE* alleles have been studied. However, due to the variable prevalence of *APOE* genotypes within human populations, characterization of cell-type specific transcriptomes across *APOE* genotypes has been challenging. Here, we integrated previous and newly generated single-nuclei sequencing (snRNA-seq) data in the prefrontal cortex (PFC) from individuals across *APOE* genotypes (*2/2*, *2/3*, *3/3*, *3/4*, *4/4*). Clustering analysis revealed distinct excitatory and microglial subpopulations that were uniquely enriched or depleted for *APOE4/4* AD. Notably, an excitatory neuronal cluster exhibited neurofibrillary tangle (NFT) signatures and was selectively depleted in *APOE4/4* AD cases. In addition, several microglial subpopulations were influenced by both *APOE4* dosage and disease status. Among these, the putative AD risk gene *FRMD4A* emerged as *APOE4* dose and AD-dependent. These findings were validated by RNAscope in an extended cohort. Together, our findings provide insights into how *APOE4* reshapes cellular states and contributes to cell-type-specific vulnerability in AD.

## INTRODUCTION

*APOE4* (*e4*) is the strongest genetic risk factor for late onset Alzheimer’s Disease (LOAD)^1^. As an estimate, having a single *e4* allele increases AD risk 2- to 4-fold, and having two *APOE4* alleles approximately 8- to 16-fold^2^. In the central nervous system (CNS), *APOE* functions as a high-density lipoprotein responsible for cholesterol and lipid homeostasis^3^. *APOE3* (*e3*), the most common isoform, is considered the parent form and is associated with normal plasma lipid levels^4^. Additionally, *APOE* regulates synaptic activity^5^, tau^6–11^, amyloid beta (Aβ)^12–14^ pathology, and debris clearance^15–17^, as well as other more cell-type specific roles^18^ ^19^. Although *e4* carriers have more heterogeneous cognitive and pathological manifestations of AD^20^, in general, *APOE4* carriers develop AD pathologies more intensely and earlier compared to non-carriers^1^ ^21–23^. The hallmarks of LOAD, encompassing amyloid-β (Aβ) pathology^24–29^, tau pathology^6^ ^30^ neurodegeneration^6^ ^9^ ^10^, and neuroinflammation^31–33^ have all been correlated with the *e4* allele (reviewed by Fernández Calle et al.^34^ and Tzioras et al.^35^). While the presence of the *e4* allele increases the risk of developing AD, *APOE*2 (*e2*) has been shown to have protective effects^36^. An allelic serial pattern of *e2* < *e3* < *e4* has been observed across several variables from lipid metabolism^37^ to transcriptomic landscapes^38^. However, human studies comparing the gene expression of disease versus control patients across *APOE* genotypes has proven difficult due to a number of factors including the limited availability of certain post-mortem tissues and low frequency of particular genotypes across populations^2^ ^39–41^. Fortunately, recent focus on developing more effective tools for data integration^42^, in combination with initiatives to make data more freely available to researchers^43–47^ has allowed for unprecedented access to a wide array of data. To this end, we utilized two publicly available snRNA-seq datasets (Mathys et al. 2019^48^ and Zhou et al. 2020^49^) and integrated newly generated snRNA-seq data from individuals with *APOE2/2* genotype and *APOE4/4* controls, enabling a direct comparison of the transcriptomic landscape across *APOE* genotypes.

APOE isoforms have cell-type specific effects, and the major brain cell types contribute uniquely to AD pathology^10^ ^33^ ^50–54^. We performed clustering analysis to determine the composition of cell types in prefrontal cortex (PFC) tissue across *APOE* genotypes (*2/3*, *3/3*, *3/4*, and *4/4*) using our integrated data set. By comparing our dataset to transcriptomic profiles of cells bearing neurofibrillary tangles (NFTs)^55^, composed of abnormally hyperphosphorylated tau, we identified a cluster of excitatory neurons (Ex5) characterized by an NFT signature. Interestingly, this neuronal subgroup was depleted in *APOE4/4* AD cases. Subclustering analysis of excitatory neurons further revealed an *APOE4*-dose dependent upregulation of immune responses and genes involved in ion regulation. We also report microglial phenotypes associated with the *APOE4* allele. Notably, we have identified a novel target, *FRMD4A*, that is overexpressed in *APOE4/4* individuals compared to controls and shows a dose-dependent increase across AD *APOE* genotypes. These findings reveal specific cell populations in both *APOE 4/4* control and AD brains, increasing our understanding of cellular vulnerability and resilience in relation to *APOE* genotype.

## METHODS

### Human postmortem brain tissues

Post-mortem human brain specimens were obtained from both the Mount Sinai/ JJ Peters VA Medical Center NIH Brain, Tissue Repository (NBTR) and the Banner Sun Health Research Institute’s Brain and Body Donation Program (BBDP)^56^, the University of Pennsylvania’s Alzheimer’s Disease Research Center (ADRC), and the Emory Goizueta ADRC/ Center for Neurodegenerative Disease (CND). For snRNA-seq, frozen post-mortem samples were obtained from the prefrontal cortex (PFC) in Brodmann Area 10 (BA10) from patients with various *APOE* genotypes. For RNAscope validation, formalin-fixed, paraffin-embedded (FFPE) samples from the same brain region (BA10) were used.

### Neuropathological criteria for grouping cohorts

For sequenced cases, individuals were separated into AD and CTRL groups according to the neuropathological assessment guidelines provided by the National Institute on Aging – Alzheimer’s Association^57^. AD neuropathological changes (“ABC”) equating to “Not” or “Low” were grouped as control (CTRL), while “ABC” scores of “Intermediate” or “High” were grouped as AD. Demographical, neuropathological, and clinical data for each case is included in **Supplemental File 1**. Due to the limited availability of tissue at the time of sequencing from *APOE2/2* patients meeting the neuropathological and clinical criteria for AD, all sequenced *APOE2/2* cases were grouped together for analysis irrespective of disease status. In total, the MSSM snRNA-seq cohort included 14 cases. These included PFC (BA10) from 7 *APOE4/4* samples (n=2 CTRL; n=5 AD) and 7 *APOE2/2* samples (not grouped by disease status). For external sequencing data from the ROSMAP cohort (Mathys^48^ and Zhou^49^), cases were grouped into AD and CTRL according to the neuropathological and clinical criteria described by the authors. A total of 40 AD, 37 CTRL, and 7 undefined cases were included in the integrated snRNA-seq dataset.

For RNAscope validation, FFPE tissue was obtained for an extended cohort of *APOE2/2*, *APOE3/3*, *APOE3/4*, and *APOE4/4* genotypes. Cases were grouped into AD or CTRL using the same neuropathological criteria as our MSSM sequencing cohort.

### Sample Preparation

For single-nuclei RNA sequencing (snRNA-seq), fresh frozen tissue was sent to Azenta Life Sciences for sample preparation. To prepare single cell suspensions, myelin debris was removed (Myelin Removal Beads II; Miltenyi Biotec; 130-096-433) and samples were passed through a column (LS Columns; Miltenyi Biotec; 130-042-401). Samples were dissociated using the gentleMACS Program (Miltenyi Biotec) in Nuclei Extraction Buffer (Miltenyi Biotec; 130-128-024) with RNase inhibitor to generate single nuclei suspensions according to the manufacturer’s protocol. Library preparation and droplet-based snRNA-seq was performed by Azenta Life Sciences using the 10x Genomics Chromium workflow.

For RNAscope *in situ* hybridization, formalin-fixed-paraffin-embedded (FFPE) tissue (BA10) blocks were cut to 5 microns under RNase-free conditions. RNAscope *in situ* hybridization (ACD, RNA-ISH) was performed, according to the manufacturer’s protocol, using the chromogenic RNAscope 2.5 LS Duplex (ACD; 322500) and the Leica Biosystems’ BOND RX Research Advance Staining System. H&E was used for cell detection. Slides were imaged at 40x using the NanoZoomer S210.

### Data processing

We identified three single-nuclei RNA-sequencing cohorts of AD and control samples with annotated *APOE* genotypes: MSSM (internal cohort), ROSMAP from Mathys et al. 2019^48^, and cohort from Zhou et al. 2020^49^. We utilized Seurat workflow^58^ ^59^ to identify quality cells with mitochondrial rate < 10% and 20 < number of expressed genes < 10,000. In addition, high-confidence doublet cells as inferred by DoubletFinder^60^ were excluded. Then, we performed integration of snRNA-seq across all samples in the three cohorts by Robust Integration of Single-Cell RNA-seq data (RISC)^61^. Briefly, RISC searches for a reference data set onto which other data sets are integrated by aligning gene-eigenvectors^5^, yielding the integrated eigen-vectors. In tandem, RISC generates batch-adjusted log-normalized expression data for further down-stream analyses. By following the recommended RISC workflow, we first evaluated which sample would serve as the most adequate reference by *InPlot()* function evaluating the qualities of gene-eigenvectors as the references to reflect as much information. These evaluations reflect (1) how many eigenvectors in each data set can explain 95% gene expression variance, indicating the number of eigenvectors to be used in the data integration, (2) which data set contains the most diverse cell population (cluster number score), (3) in which data set gene expression variance would be explained by more eigenvectors (principal component score), and (4) whether the eigenvectors in each data set are normally distributed (Kolmogorov-Smirnov score) (**Supplemental Figure 1**). Per recommendations from the authors, we selected the best scoring sample in Cluster Num. score as the reference (marked as Set 6 in **Supplemental Figure 1a-c**). Then, the samples were integrated using *scMultiIntegrate()* with 15 eigen-vectors. The integrated eigen-vectors were utilized as the Principal Components (PCs) from which the respective UMAP embeddings were calculated. We then performed Louvain clustering with resolution at 0.5 by utilizing k-nearest neighbor clustering implemented in Seurat^58^.

### Comparative analysis between inhibitory and excitatory neurons

We hypothesized that excitatory neurons (Ex) are more vulnerable towards neurofibrillary tangles (NFTs) than inhibitory neurons (In). Meanwhile, Otero-Garcia *et al*. 2022 identified molecular signatures of NFT-bearing neurons through single somas of NFTs^55^. We utilized these NFT signatures and compared their differential expressions between Ex and In neurons. The differential expression analysis was performed through generating cell type-specific pseudo-bulk data by aggregating cells from the same samples within each cell type. Then, we applied DESeq2^62^ framework on the pseudo-bulk data with the model, y ∼ cohort + gender + disease (AD/control) + cell type (Ex/In). The differentially expressed genes (DEGs) were then identified by FDR < 0.05, fold change > 1.2 or < 1/1.2 (**Supplemental File 3**).

### Murine NFT signature

QC was accomplished using Seurat with the criteria of mitochondrial rate < 25% and 500 < # of genes < 6,000, and removal of high confidence doublets using DoubletFinder^63^. *AddModuleScore()*^64^ was used to calculate overall expressions of the human cell type marker genes in the murine clusters. (**Supplemental Figure 4)**.

### Excitatory neuronal subclustering

Based on the clustering results on the full data set, we extracted the excitatory neuron clusters (Ex1-5) and performed subclustering on these cells. Firstly, we evaluated the variances of the batch-corrected RISC eigenvectors to identify variable RISC eigenvectors within the excitatory neurons, and identified 1st, 3rd, 5th, 11th and 12th eigenvectors show relatively higher variability amongst the excitatory neurons (**Supplemental Figure 1d**). We utilized these variable RISC eigenvectors to construct the shared nearest neighbors by *FindNeighbors()* function, then performed clustering by Louvain clustering with γ=0.8 by *Findclusters()* function in Seurat R package (v 5.0.0). These yielded a total of 9 subclusters (**Figure 2c**).

**Figure 1.**
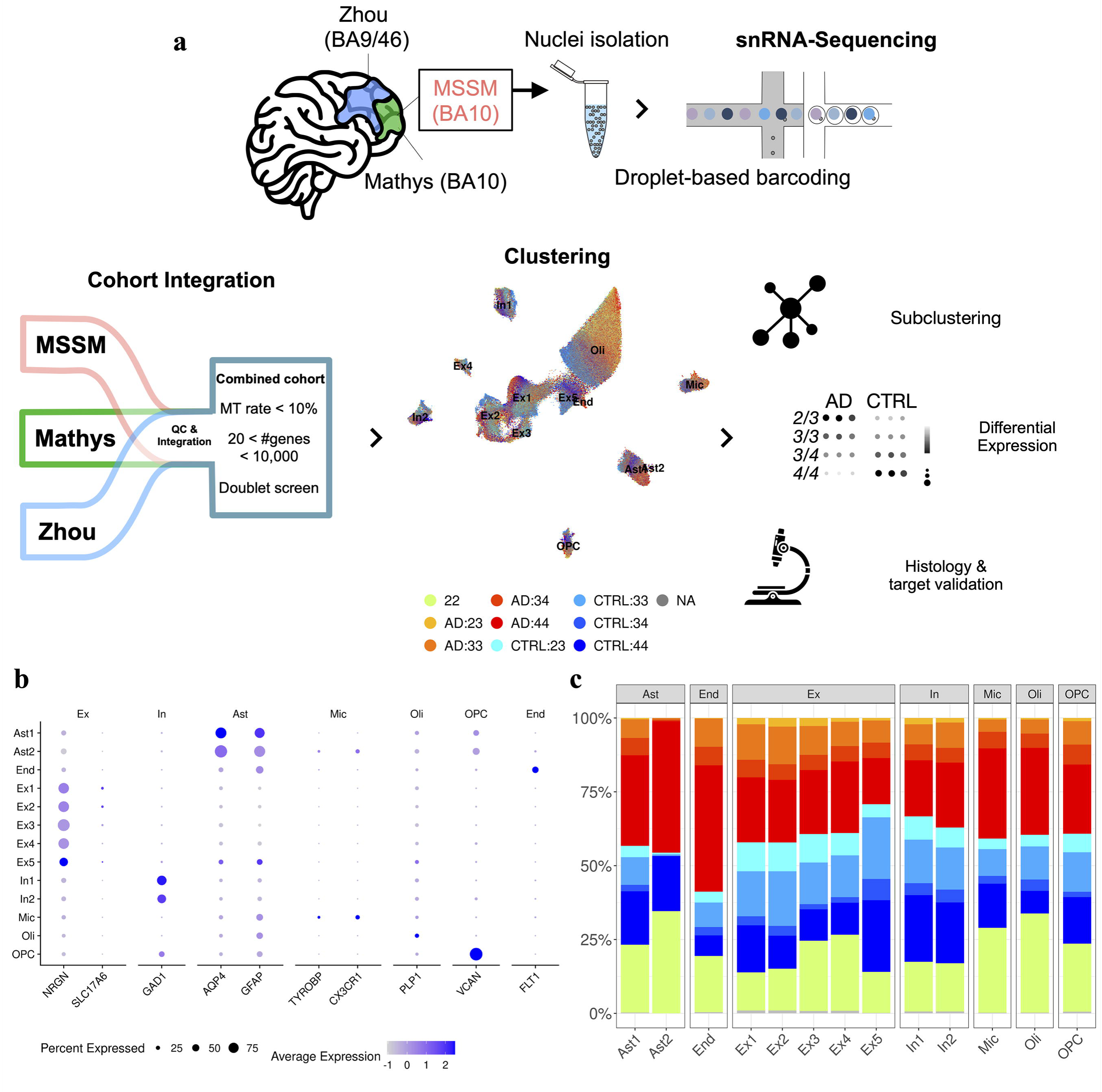
a) Schematic overview of integrated cohort. Frozen post-mortem prefrontal cortex (PFC – Brodmann Area shown) tissue from three unique cohorts – Mathys^48^, Zhou^49^, and MSSM (this study) was subject to nuclei isolation followed by droplet based snRNA-sequencing. This data from all three cohorts was integrated by Robust Integration of scRNA-seq data (RISC). Each cohort was processed with quality control (QC) filters described. UMAP plot showing the major cell populations of the integrated snRNA-seq transcriptome landscape. Pathological AD or control (CTRL) patients and APOE genotypes are shown in the legend. Clustering, subclustering, and differential expression analysis was performed using the integrated data set, followed by histology and target validation. b) Cell type marker expression dot plot in different clusters. c) Proportion of cells coming from different AD/CTRL cells with different APOE genotypes.

**Figure 2.**
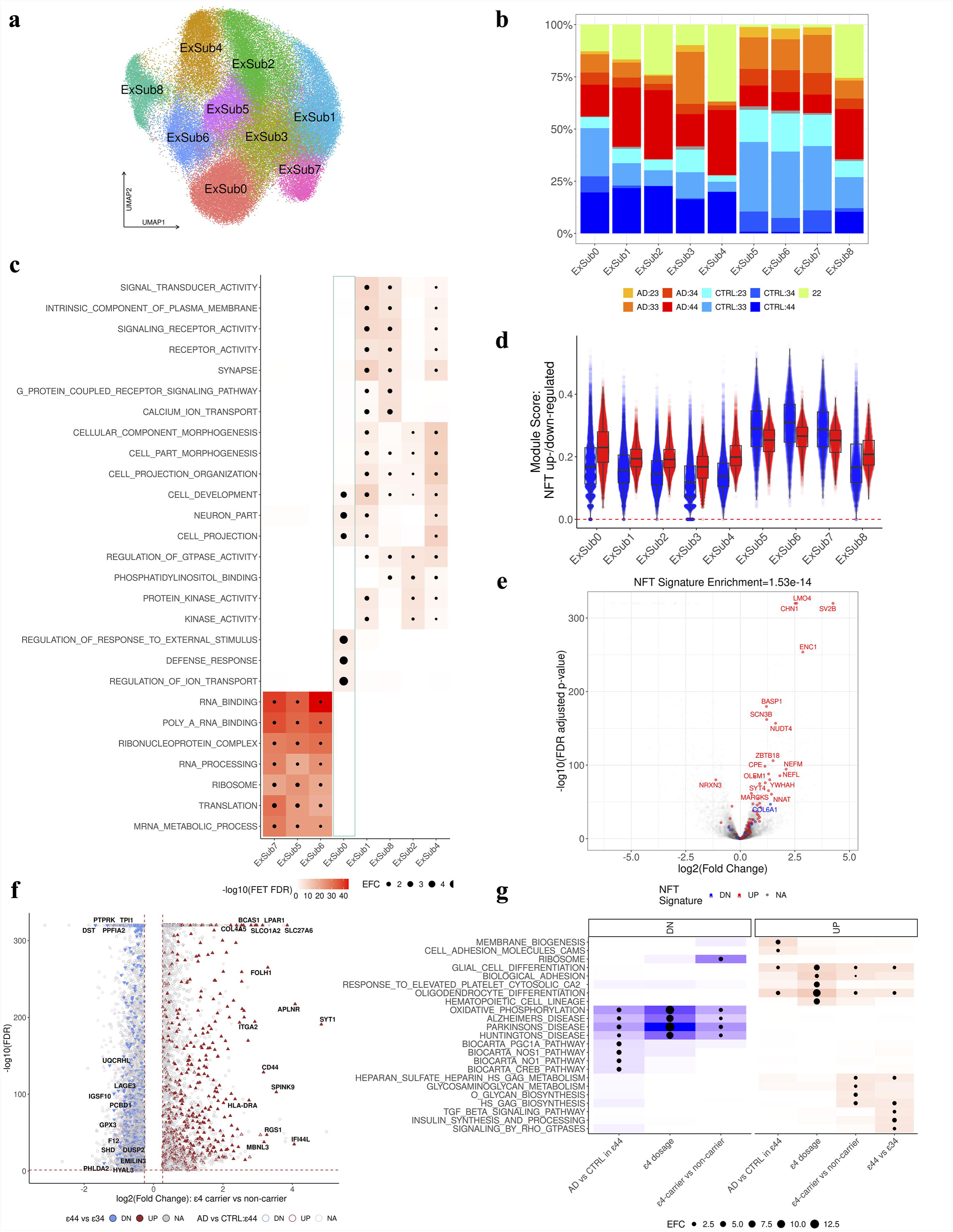
NFT-signature bearing excitatory neuronal cluster 5 is comprised of subclusters that differ based off APOE genotype. a) UMAP plots of excitatory neuron subclusters by APOE genotype. b) Barplot shows proportion of different cell groups by APOE genotype in different subclusters. The cell groups are shown in the legend below. c) Heatmap of significantly enriched pathways (FDR-adjusted Fisher’s Exact Test (FET) *p*-value < 0.05) in the subcluster markers. Collections of top 4 enriched pathways in different subcluster markers are shown. The dot size is proportional to the enrichment fold change. d) Cellwise module scores of NFT signatures (y-axis) in different excitatory neuronal subclusters (x-axis). Red: module scores of upregulated NFT signature, Blue: module scores of down-regulated NFT signature. e) Volcano plot of differentially expressed genes in excitatory neurons against inhibitory neurons. X-axis: log2(fold change), Y-axis: -log10(FDR). NFT signatures are highlighted as red (upregulated) and blue (downregulated) with upregulated NFT enrichment p-value in upregulated genes in excitatory neurons on the top. f) Volcano plot (x-axis log2 fold change, y-axis: -log10(FDR) to show differentially expressed genes (DEGs) between e4 carriers and non-carriers in excitatory neurons. Significant e4 carrier DEGs (FDR < 0.05, fold change > 1.2 or < 1/1.2) that are upregulated in APOE4/4 AD compared to APOE4/4 CTRL, and upregulated APOE4/4 AD compared to APOE3/4 AD are labeled by gene symbols. g) Significantly enriched pathways in gene signatures of e4 dosage and differentially expressed genes in AD are shown.

### Microglial subclustering

Upon subsetting the microglia cells from the initial unsupervised clustering results across the three cohorts (MSSM, Mathys and Zhou), we selected informative RISC eigenvectors^61^ as the clustering features for the microglial cells by comparing their mean values across the microglial cells, and applying > 2E-5 as the threshold for the feature selection (**Supplemental Figure 1e**). We then performed the shared nearest neighboring approach implemented in Seurat to identify the clusters with the resolution at 0.8^58^. This yielded 11 microglial subclusters (**Figure 3a**). The subcluster markers within the microglial cells were then identified by *FindMarkers()* function in Seurat via MAST method^65^ with FDR < 0.05 and fold change > 1.

**Figure 3.**
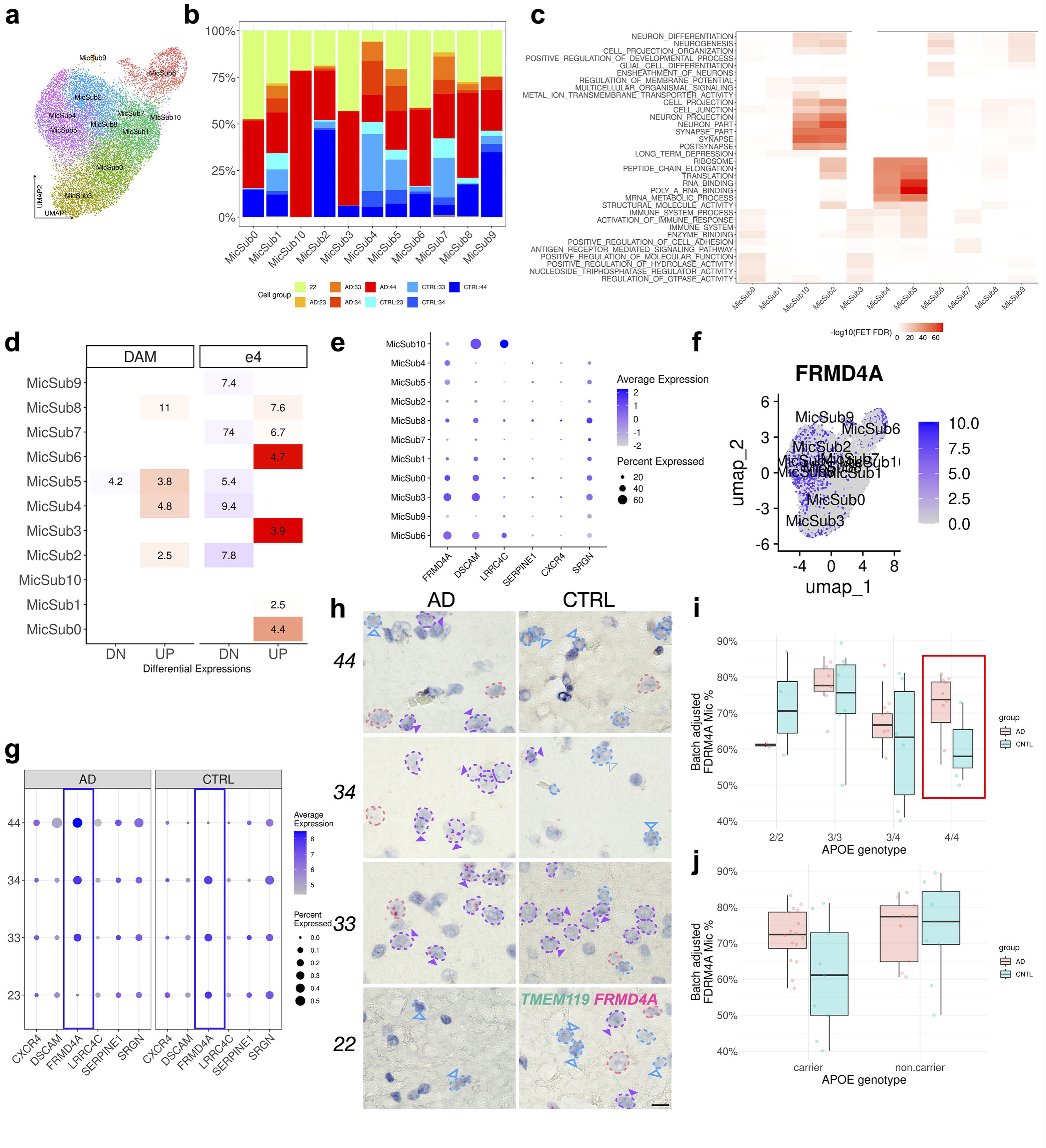
Microglia display e4-specific transcriptional states. a) UMAP plot of microglial subclusters. b) Proportions of cells from subjects with different APOE genotypes in microglial subclusters. c) MSigDB pathway/function enrichments in microglia subclusters. Collection of top 5 pathways/functions from different subclusters are shown. d) Enrichments of disease-associated microglia (DAM) signatures (left) and e4 dosage signatures (right) in microglial subclusters. X-axis: up /down-regulated in each signature category, y-axis: microglia subcluster markers. Heatmap colors correspond to log10(Fisher’s Exact Test FDR). e,f) Dot plot (e) and feature plot (f) to show expressions of top e4 dosage associated genes in microglia subclusters. g) Dot plot of the top 9 APOE-dose dependent genes expressed in microglia. An allele-specific, dose-dependent pattern of FRMD4A expression that is dependent upon AD pathology based off the following criteria: differentially expressed between AD e4 carriers and AD non-e4 carriers, and differentially expressed between AD APOE4/4 and AD APOE3/4 cells h) Representative images of H&E stained PFC tissue stained with RNAscope probes for FRMD4A (magenta) and TMEM119 (green). Scale bar = 10 µm. i,j) Bar plots showing FRMD4A expression in microglia (percent of FRMD4A+;TMEM119+ cells/total TMEM119+ cells) from AD and CNTL cases by (i) APOE genotypes and (j) non e4 carrier versus e4 carrier.

### *e4* dose-dependent genes in microglial cluster 5

The *e4*-dosage dependent gene expressions were then evaluated with the following criteria: 1) Significantly upregulated genes in AD *e4* carriers, compared to AD non-*e4* carriers: The significant up-regulations were identified with FDR < 0.05, and fold change > 1.2. Among the significantly upregulated genes, we also enforced < 10% of non-*e4* carrier cells to express them to ensure *e4*-carrier specificity. 2) Upregulated genes in *APOE4/4* AD compared to *APOE3/4* AD: Among the significantly up-regulated genes in AD *e4*-carrier cells, we further identified significantly upregulated genes in the AD homozygotes (*APOE4/4*), compared to the AD heterozygotes (*APOE3/4*) with FDR < 0.05 and fold change > 1.2.

### Quantitative assessment of *FRMD4A* expression with RNAscope

To quantify the average number of *FRMD4A* spots per total cells, one probe targeting human *FRMD4A* (ACD; 1266448-C1, green) was used and *FRMD4A* was quantified in all cell types in cortical layers II/III using QuPath and ImageJ. To quantify the number of *FRMD4A* positive microglia, two probes, *FRMD4A* (ACD; 1266448-C2, red) and *TMEM119* (ACD; 478918, green) were co-stained and *FRMD4A* expression was quantified manually by a blinded researcher. Approximately 150 individual cells from layers II/III were classified as *FRMD4A*+, *TMEM119*+, or dual *FRMD4A*+/*TMEM119*+. The number of *FRMD4A+/TMEM119+* (dual positive) cells over the number of *FRMD4A+/TMEM119+* (dual positive) plus *TMEM119+* (microglial marker only) cells was calculated to determine the number of *FRMD4A+* microglia of all microglia counted (percent *FRMD4A+* microglia). These values were then corrected for batch number (run number) and for brain bank source using linear regression model. We utilized *removeBatchEffect()* function in limma R package (v 3.54.2) to model the read out values, y, by y ∼ experiment batch + biobank source + *APOE* genotype + disease status (AD/Control) + error. The read out values were adjusted for experiment batch and biobank source variables.

### Differential Expression Analysis by Cell Type

Differential expression (DE) analyses were conducted across several cell types, including astrocytes (Ast), oligodendrocytes (Oli), excitatory neurons (Ex), inhibitory neurons (In), oligodendrocyte precursor cells (OPC), microglia (Mic), and endothelial cells (End). Each analysis was performed independently for each cell type and *APOE* genotype. Genes were considered upregulated if they had a log (fold change) greater than 1.2 and an adjusted *p*-value less than 0.05. Conversely, genes were considered downregulated if they had a log (fold change) less than −1.2 and an adjusted *p*-value below 0.05. Then, overlapping and unique DEGs were identified across all possible pairwise combinations where DEG sets within that cell type were identified. A list of these genes can be found in **Supplemental File 3.**

### Gene Ontology Enrichment by Cell Type

Pathway enrichment analysis was performed for each set of differentially expressed genes using the Molecular Signature Database (MSigDB) v6.0^66^ and GOtest R packages. *GOtests* applies a one-sided hypergeometric test to assess the statistical significance of overlap between the DEG sets and curated gene sets within Gene Ontology (GO) categories. A complete list of enriched pathways can be found in **Supplemental File 4.**

## RESULTS

### Single nucleus characterization of *APOE* gene expression across multiple cohorts

To identify *APOE4*-associated molecular signatures of vulnerability in AD, we performed snRNA-seq on human postmortem PFC (BA 10) tissues from AD *APOE4/4* (*n*=5) and corresponding *APOE4/4* controls (*n*=2). snRNA-seq was also performed on PFC tissue from *APOE2/2* individuals with varying neuropathological and clinical characteristics (*n*=7). Clinical and neuropathological demographics for our cohort (MSSM) are summarized in **Supplemental File 1**. To enhance our analytical power, we included *APOE3* genotypes (*2/3*, *3/3*, and *3/4*) by integrating our MSSM dataset with two publicly available datasets: ROSMAP from Mathys *et al.* 2019^48^ and Zhou *et al.* 2020^49^ (**Figure 1a**). A total of 40 AD, 37 CTRL, and 7 undefined cases were included in the integrated snRNA-seq dataset.

For total sample composition of the integrated dataset, see **Supplemental File 2.** Taken together, we profiled a total of 333,648 single-nucleus transcriptomes (**Supplemental File 2).**

### Cell Type Composition and Cluster Enrichments Depend Upon *APOE* Genotypes and Disease State

The single-nuclei transcriptomic data captured all major brain cell types. We performed graph-based clustering and visualized the single-nuclei transcriptomes in the uniform manifold approximation and projection (UMAP) space, which revealed 13 distinct cell clusters (**Figure 1a**). Clusters were annotated based on canonical gene expression of known cell type markers: excitatory neurons (*NRGN* and *SLC17A6*), inhibitory neurons (*GAD1*), astrocytes (*AQP4* and *GFAP*), microglia (*TYROBP* and *CX3CR1*), oligodendrocytes (*PLP1*), oligodendrocyte progenitor cells (OPCs) (*VCAN*), and endothelial cells (*FLT1*) (**Figure 1b).**

We hypothesized that certain cell populations would be enriched in the *APOE4/4* AD group. To this end, we tested enrichment of different APOE genotypes in the cell clusters by Fisher’s Exact Test (FET) (**Figure 1c**; **Supplemental Figure 2d**). Indeed, the cell clusters showed heterogeneous distributions of different APOE genotypes, and we observed robust over-representations of *APOE4* alleles in AD amongst microglia, endothelial cells, astrocytes and several excitatory neuron clusters (Ex1, 2). On the contrary, Ex5 showed relative depletion of cells with *APOE4* alleles.

#### Excitatory neuronal subclustering

To dissect the heterogeneity of APOE genotypes amongst the excitatory neurons at a greater resolution, we performed subclustering analysis within the excitatory neurons and identified nine distinct subclusters across all cohorts (**Figure 2a, Supplemental Figure 3**). Differentially expressed genes (DEGs) were identified for each subcluster (**Supplemental File 4**), providing insights into differences in cell-type proportions between AD and controls across *APOE* genotypes (**Figure 2b**), as well as enriched biological pathways within each subcluster (**Figure 2c, Supplemental File 5**).

ExSub0 and ExSub8 exhibited distinct transcriptional profiles. When we compared the proportion of groups making up each excitatory subcluster, we found that ExSub0, the homologous subcluster to Ex5, showed robust depletion in AD *APOE4/4* post-mortem brains while ExSub8 was depleted in *e4* carrier controls (**Figure 2b**). Identifying the transcriptional signatures of ExSub0 and ExSub8 may shed insight into what drives differences in cellular responses from individuals with the same genotype (i.e. *APOE4/4*) resulting in differential disease states. Therefore, we focused on significant genes dominating ExSub0 and ExSub8.

The three top enriched pathways for ExSub0 were Regulation of Response to External Stimulus (*AGT*, *MIF*, *MT3*), Regulation of Ion Transport (*ATP1B2*, *CALM2*), and Neuron Part (*SNCG*, *TUBB3*) (**Supplemental File 6**). Another top enrichment, which was unique for ExSub0, was Defense Response (*B2M*, *COX8A*, *SPP1*, *F3*). Further, ExSub0 was enriched for genes associated with astrocyte-derived neurogenesis including: *GFAP*, *APOE*, *AQP4*, *GAD1*, and *VIM*^67^ ^68^ and autophagy-related genes *HSPA8*, *TUBB2B,* and *GABARAP*. Following injury, cortical astrocytes have been shown to elicit a reprogramming response toward a neuronal stem cell state, eventually giving rise to neurons^67^. The neurogenic potential of GFAP-expressing cells has also been observed in the aging dentate gyrus^69^. Our findings suggest that ExSub0 may represent a neurogenesis-associated population whose depletion in AD *APOE4/4* cases could reflect impaired regenerative capacity in this genotype.

For ExSub8, which we found to be depleted in e4 carrier controls (**Figure 2b**), the top enriched pathways were Signaling Receptor Activity (*RGMA*, *ACSL1*, *GRIK4*), Calcium Ion Transport (*RYR3*, *MCU*, *CAMK2D*, *CAV1*, *ATP2B4*, *CASK*) and Receptor Activity (*MCC*, *DPP4*, *CDHR3*). Selectively vulnerable neurons characterized by Leng et al.^70^ exhibited substantial overlap (85 genes) with our ExSub8. The shared genes of these two potentially vulnerable neuronal subpopulations include canonical WNT signaling genes (*TLE4*, *CDH11*, *PCDH11Y*, *PLCB4*, *CDH18*), Phosphoinositide signaling (*PLCB4*, *DGKG*, *DGKD*), and additional ion homeostasis genes (*TRPC1*, *TRPC6*, *SLC24A3*, *SLC4A4*, *KCNIP1*).

#### An Excitatory Neuronal Subcluster Is Enriched for Tau Signatures

Otero-Garcia et al.^55^ developed a method for profiling NFT-bearing neurons from the human AD brain. NFT-bearing neurons characteristically upregulate certain genes involved in synaptic transmission, oxidative phosphorylation, and mitochondrial dysfunction^55^. We compared the signatures of the excitatory neuronal populations from the MSSM, Mathys, and Zhou cohorts with murine NFT-signatures identified by Otero-Garcia et al.^55^ (**Figure 2d, Supplemental Figure 4a,c).** Subcluster ExSub0 overlapped significantly with NFT-signatures (**Figure 2d, Supplemental Figure 4e**). Additionally, ExSub5, 6, and 7 had high module scores for NFT- up- and downregulated genes (**Figure 2d**). As previously mentioned, enriched pathways of ExSub5, 6, and 7 were dominated by RNA regulatory pathways, while ExSub0 was primary associated with defense response (**Figure 2c**). The enrichment of RNA regulatory pathways in ExSub 5, 6, and 7 could be explained by increased accumulation of RNA in NFTs^72,73^. Next, we identified DEGs enriched in excitatory versus inhibitory neurons that overlapped with NFT-signature genes (**Figure 2e, Supplemental File 7).** Among these, commonly upregulated genes were cytoskeletal genes (*ENC1*, *BASP1*, *NEFM*, *NEFL*), calcium signaling (*LMO4*, *CAMK2N1, CALM1*), genes involved in signal transduction (*CHN1*, *YWHAH*), axonal growth (*OLFM1*, *CPE*), ion channels (*SCN3B*, *NNAT*), and vesicle secretion (*SYT4*, *SV2B*, *BASP1, MARCKS*). For example, *OLFM1*, an NFT-signature gene also upregulated in our cluster Ex5, has been shown to regulate axonal growth ^71^ ^72^ and interacts with synaptic proteins^73^. Brain-abundant membrane attached signal protein 1 (BASP1) controls the movement of cytoskeletal-related proteins, thereby enhancing axon growth and plays important roles in neurite outgrowth, both during development and in adulthood^74^ ^75^. *BASP1* and *MARCKS*, which were both upregulated in Ex5, belong to a family of growth-associated proteins which locally alter their gene expression in axon terminals^76^. Another upregulated NFT gene, *CPE* or carboxypeptidase E, has a dual role: exhibiting both exopeptidase activity, as well as acting as a secretory sorting receptor^77^ ^78^. CPE has previously been implicated in both MCI and AD^79^ ^80^ and accumulates in senile plaques of AD patients ^81^. Interestingly, CPE may act as a neurotrophic factor, upregulating levels of pro-survival protein BCL2^82–85^. Similar to the findings presented by Otero-Garcia, these NFT-signature genes are not dominated by cell death pathways.

We compared Ex5, homologous to subcluster ExSub0, against snRNA-seq data from a tau (P301S) *APOE4* mouse model previously generated by Wang *et al.*^10^ (**Supplemental Figure 4**). In this study, removal of astrocytic *APOE3* and *APOE4* expression was accomplished by *Aldh1l1*-driven tamoxifen-inducible Cre expression to produce TAFE3 and TAFE4 mice (Tau/*Aldh1*∣*1*-CreERT2/*Apoe3^flox/flox^* and Tau/*Aldh1*∣*1*-CreERT2/*Apoe4^flox/flox^*)^10^. At 9.5 months-of-age, hippocampal lysates were harvested from TAFE3 and TAFE4 mice and compared to age-matched controls injected with oil (vehicle). Here, transcriptomic data from oil-treated *APOE4* floxed-mice was used as a murine counterpart to identify similarities in tau pathology in excitatory neurons. All major brain cell types were identified in the murine data (**Supplemental Figure 4d,e**). An excitatory neuron subpopulation Cluster 10 (Ex10) in the murine snRNA-seq data has a marker signature similar to that of Cluster 5 in the human snRNA-seq data (Wilcoxon rank-sum *p*-value = 1.03E-165, 1.35-fold increase; **Supplemental Figure 4f**) and the human NFT signature (Wilcoxon rank-sum *p*-value = 2.33E-69, 1.4-fold increase; **Supplemental Figure 4g**). Similar transcriptional signatures could be seen between Ex5 and NFT-signature associated murine Ex10, where *Nrgn*-expressing excitatory neurons expressed *Aqp4* and *Gfap* (**Supplemental File 7**), indicating a conserved NFT-signature that exists between mouse and human. Interestingly, this NFT-signature significantly overlaps with downregulated genes in AD versus CTRL *APOE4/4* cases (**Supplemental Figure 4c**). Moreover, there is a substantial depletion of Ex5 and its homologous subcluster, ExSub0, in *APOE4/4* AD cases compared to other AD genotypes, suggesting a genotype-specific vulnerability (**Figure 1c** and **Figure 2b**). This suggests that loss of this subpopulation of excitatory neurons, which may be involved in defense response and neurogenesis, may contribute to the vulnerability of excitatory neurons in *APOE4/4* cases.

#### Excitatory Neurons Exhibit an APOE4-Dosage Dependent Signature

To further explore the impact that e4 dosage has on the gene expression of excitatory neurons, we selected DEGs that satisfied the following criterion: 1) differentially expressed between *e4* carriers versus non-carriers, 2) differentially expressed between AD *4/4* patients versus AD *3/4* patients, and 3) differentially expressed between AD and control *APOE4/4* patients in the same direction. Genes meeting these criteria, referred to as *APOE*-dosage dependent DEGs, are listed in **Supplemental File 8**. 353 upregulated *APOE*-dosage dependent genes were found in excitatory neurons (**Figure 2f**); this list included genes potentially involved in antiviral immune response (*SPP1*^86^, *ZDHHC11*, *ZDHHC11B*^87^ ^88^, and IFI44L^89^), *LAMP2*, a keystone autophagy regulator, *OXR1*, an oxidative stress gene^90^ ^91^, the guanine-nucleotide exchange factor (GEF) triggered by inflammation, *RASGEF1B*^92^ and several genes involved in ion regulation (*PIEZO2*^93^, *KCNQ5*^94^, *S100A16*^95^, *CLCA4*^96^). Of note, Mathys et al., 2019 described *RASGEF1B* as a marker for an excitatory neuronal subpopulation associated with AD pathology^48^. Here, we were able to further define *RASGEF1B* as an *APOE4* dose dependent gene. The pathways that dominated the upregulated genes included Response to Elevated Platelet Cytosolic Ca^2+^, Hallmark Coagulation, and Cell Surface (**Figure 2g, Supplemental File 9**). 456 genes were found to be significantly downregulated in an *APOE*-dosage dependent manner. These included: *DST*^97^, a cytoskeletal gene, *SIK3*, a positive regulator of mTOR^98^, and *CDH18*, a cadherin, and *CAMKK2*, a calcium-dependent kinase, and *GPX3*, a selenoprotein which induces mitochondrial-mediated apoptosis^99^ ^100^ (**Figure 2f**).

In general, the *APOE4* dose-dependent molecular signature was markedly distinct from NFT-signatures. In fact, there was a significant overlap between *APOE4*-dosage downregulated genes and NFT signature upregulated genes (**Supplemental Figure 4c**). This supports the idea that other pathologies, outside of NFT accumulation, may also drive the detrimental effects of *APOE4* on neurodegeneration ^101–103^. An alternative possibility is that this represents an NFT-neuronal subpopulation that was more resistant to death while others may have disappeared.

### Microglia show preferential expression in *APOE4/4* compared to other *APOE* genotypes

To assess whether specific *APOE* genotypes were enriched for particular cell types, we examined genotype-specific cluster distributions in AD and control across all cohorts (**Supplemental Figure 2**). Enrichment for microglia was seen in *e4* carriers in the Mathys cohort (**Supplemental Figure 2d**). Among AD cases, *APOE4* carriers showed significant enrichment of microglia compared to non-carriers in all three cohorts (**Supplemental Figure 2d**). Control *APOE4* carriers also showed a significantly greater proportion of microglia in the MSSM and Mathys cohorts but not in the Zhou cohort, although these groups were limited in sample number (**Supplemental Figure 2d**).

#### Microglial subclustering reveals distinct e4 microglial signature

Microglia exist in dynamic states that can be indicative of and contribute to the overall condition of surrounding cell types. Microglial subclustering yielded 11 distinct subclusters (**Figure 3a, Supplemental Figure 5**) and revealed heterogeneous distributions of cells with different *APOE* genotypes (**Figure 3b**). Particularly, the subcluster markers highlighted distinct DEGs and pathways underlying them (**Figure 3c, Supplemental Files 10,11**), indicative of distinctive microglial transcriptional states. Interestingly, MicSub8, a microglial subcluster with high expression of *HIF1A* was enriched for *APOE4/4* AD cases (**Figure 3b**).

To better differentiate microglial subclusters with commonly enriched pathways, each microglial subcluster was assigned a gene, or pair of genes, that were unique to that subcluster and best represented the identity of the entire set of significantly enriched genes for that subcluster. MicSub0 (*P2RY12+;MEF2A+*) and MicSub3 (*P2RY12+; CX3CR1+*) shared a set of common enriched pathways including immune-related pathways (Immune System Process, Activation of Immune Response, and Immune System) and GTPases (Positive Regulation of Hydrolase Activity, Nucleoside Triphosphatase Regulator Activity, and Regulation of GTPase Activity) (**Supplemental File 11**). Both subclusters also appeared more homeostatic in nature, expressing known homeostatic genes including *P2RY12*, *NFATC2*, *EP300*, *BACH1*, *IKZF1*, *TCF7L2*, and *FOXO3*^104^ ^105^. MicSub2:*CALM1+*, MicSub8:*HIF1A+;APOE+*, and MicSub10:*HS6ST3+* were commonly enriched for Neurogenesis, Neuron Projection, and RNA Binding. However, there were some patterns unique to each of these subclusters. MicSub2 was uniquely enriched for mitochondrial and oxidative pathways (Oxidative Phosphorylation, Respiratory Electron Transport, Mitochondrial Membrane). MicSub8 was uniquely enriched for Positive Regulation of Nitric Oxide Synthase Activity and Organic Acid Transport. Additional pathways enriched specifically in this subcluster included Regulation of Reactive Oxygen Species Metabolic Process, Establishment of Mitochondrion Localization and Neuron Projection Regeneration. The top marker gene expressed by MicSub8 was the transcription factor hypoxia-inducible factor (*HIF1A*), with *APOE* also ranking among most enriched genes. MicSub10 was uniquely enriched for Divalent Inorganic Cation Transmembrane Transporter Activity, Neuroactive Ligand Receptor Interaction, and Calcium Ion Transmembrane Transporter Activity. Microglial subclusters MicSub6:*ABCA8+* and MicSub9:*NAMPT+* were predominantly enriched for neurogenesis pathways (Neurogenesis, Neuron Differentiation, and Ensheathment of Neurons), indicating that these subclusters may represent a group of neuron protective microglia. Additional microglial subclusters included: Ribosome /Immune Response microglia (Ribosome/Immune: MicSub4:*TREM2+* & MicSub5:*GAS6+*), T-cell Receptor Pathway microglia (TCR: MicSub7:*USP39+*) and Transmembrane Transporter microglia (Transmembrane transporter: MicSub1:*CACNB4*), which may represent distinct microglial activation states. Closely related subclusters may be enriched for similar pathways (mainly inflammatory response), but notably, are enriched for distinct genes. For example, *GAS6* and *OCIAD1*, the distinct markers of MicSub5, both promote microglial phagocytosis^106^ ^107^, as does *TREM2*^108^ ^109^, which is enriched in MicSub4. However, *TREM2* may be responsive to specific inflammatory environments involving Aβ^110^ ^111^ and/or lipid dysregulation^112–114^.

Subcluster markers were enriched for murine disease-associated microglia (DAM) signature described by Keren-Shaul *et al.* 2017^104^, as well as for microglia- specific *e4* carrier signatures (**Figure 3d**). MicSub2 & 8, the synapse development associated microglia, were enriched for both DAM and *e4* carrier signatures. Microglial subclusters that were unique to e4 signatures included MicSub0, 1, 3 and 6.

A previous study identified *APOE*-linked microglial transcriptomic changes in AD following an *e2* < *e3* < *e4* expression pattern^38^. Thus, we sought to determine genes in our microglial cluster with a similar dose-dependent expression pattern. To this end, we performed differential gene expression using batch-adjusted, log-normalized expression data from RISC between AD *e4* carriers and AD non- *e4* carriers using MAST^59^ for each cell cluster. We first found significantly up-regulated genes in AD *e4* carriers compared to non-carriers, and among those genes, we then identified significantly up-regulated genes in AD *e4* homozygotes compared to AD *e4* heterozygotes. Overall, we identified 717 *e4*-dosage dependent microglial genes (**Supplemental File 8**), with the top 6 genes highlighted in **Figure 3e**.

Among the top 6 nominated targets of AD-specific *e4*-carriers, we observed that *FRMD4A* (FERM Domain-Containing Protein 4A), encoding a scaffolding protein was upregulated in MicSub0, 3 and 6 (**Figure 3f**). Enrichment of *FRMD4A* in microglia has previously been reported in neuronal surveillance subpopulations (MG1) reported by Sun *et al.* 2023^115^. Interestingly, a homeostatic microglial transcriptional state hallmarked by the expression of *FRMD4A* has been previously identified in AD and aging by Lee et al., 2023^116^.

The *FRMD4A* microglial subtype declines with age and shows a downward trend with increasing Braak stage. *ARHGAP24*, *ARHGAP22*, and *ARHGAP15*, which were highly expressed in MicSub0 and MicSub1 (*FRMD4A*-expressing microglia) are regulators of small GTPases and is involved in the actin cytoskeleton and cell polarity^117^ ^118–121^ as well as migration^122^ ^123^.

### Validation with RNAscope

Comparison of different *APOE* genotypes between AD and CTRL microglia revealed that higher *FRMD4A* expression coincides with the presence of the *e4* allele in AD, but not in CTRL (**Figure 3g**). In fact, the pattern of *FRMD4A* expression for CTRL cases exhibits the opposite pattern; where *APOE4/4* cases have the lowest average expression and percent expressed compared to the other CTRL cases.

To validate the dose-dependent expression pattern of microglial *FRMD4A*, we used RNAscope in situ hybridization in an extended cohort of post-mortem brains of AD and CTRL with *APOE2/2*, *3/3*, *3/4*, and *4/4* genotypes. Chromogenic probes targeting *FRMD4A* (red) and microglial marker *TMEM119* (green) were used, in combination with H&E staining (**Figure 3h**). The percentage of *FRMD4A* positive microglia (*TMEM119*+) was quantified (**Supplemental File 12**). Generally, the pattern of *FRMD4A* expression in microglia measured by RNAscope was consistent with snRNA-seq expression data (**Figure 3i,j**). Direct comparison of *APOE4/4* AD (74.779%±2.443 Standard Error of the Mean (SEM)) versus CTRL (58.474%±7.240) cases showed a higher number of *FRMD4A*+ microglia (**Figure 3i**). However, this did not reach statistical significance due to insufficient sample size (*p=0.142, t test*). Amongst *e4* carriers with a larger number of samples, pairwise comparison of AD (72.051%±1.875) versus CTRL (60.460%±5.083) showed a higher percentage of *FRMD4A*+ microglia in AD versus CTRL with moderate statistical significance (*p=0.0575, t test*). Overall, while no statistical significance was found between AD and CTRL groups, possibly due to inter-sample variability, the pattern of *FRMD4A*+ microglial abundance was generally higher in AD cases of *e4* carriers, supporting our snRNA-seq data.

### Differentially Expressed Genes between AD and CTRL cases are Enriched for Certain Pathways in Excitatory Neurons and Microglia

Direct gene expression comparison in AD versus CTRL groups for the most frequently occurring *APOE* genotypes within the global population^2^ were made for each major cell type (Ex, Mic, Ast, End, In, OPC, and Oil). Due to the dramatic impact of *APOE* genotype on the transcriptional signatures of excitatory neurons and microglia we had previously observed, we decided to focus on these two major cell populations for DE analysis. A full list of DEGs for all cell types can be found in **Supplementary File 3**. A full list of enriched GO terms for each intersection and across all cell types is provided in **Supplementary File 4.**

For excitatory neurons, the *APOE* genotype with the highest number of DEGs for both downregulated (**Figure 4a**, 1460 DEGs) and upregulated (**Figure 4b**, 1216 DEGs) genes was *APOE4/4*. The Venn diagrams presented in **Figure 4** highlight selected enriched pathways. Genes uniquely downregulated in the *APOE3/3* genotype show enrichment in protein translation pathways, while those specific to *APOE3/4* are associated with RNA processing (**Figure 4a**). Complementary to what we observed previously, neurogenesis was among the significantly enriched downregulated pathways in *APOE4/4* cases (**Figure 4a**). Among the downregulated neurogenesis DEGs were *PAK3*, *SOX2*, *SOX6*, *PROX1*, and *FOXG1*. Of interest, PAK3 interacts with amyloid precursor protein (APP) and regulates apoptosis^124^. Conversely, for *APOE3/3* and *APOE3/4*, neurogenesis was enriched among the upregulated DEGs (**Figure 4b**). In *APOE4/4* upregulated DEGs, ensheathment of neurons was upregulated. Disruptions in lipid metabolism caused by *APOE4* expression may give rise to dysregulated myelination of axons in *APOE4* excitatory neurons, leading to the upregulation of ensheathment-related genes (*PLLP*, *MAG*, *MYRF*). *CASP2*, which has been found to accumulate in response to proteostatic stress^125^, was among the upregulated DEGs in *APOE4/4* AD cases, along with other genes involved in neuronal death (*PARP2*, *UNC5B*, *HIPK2*).

**Figure 4.**
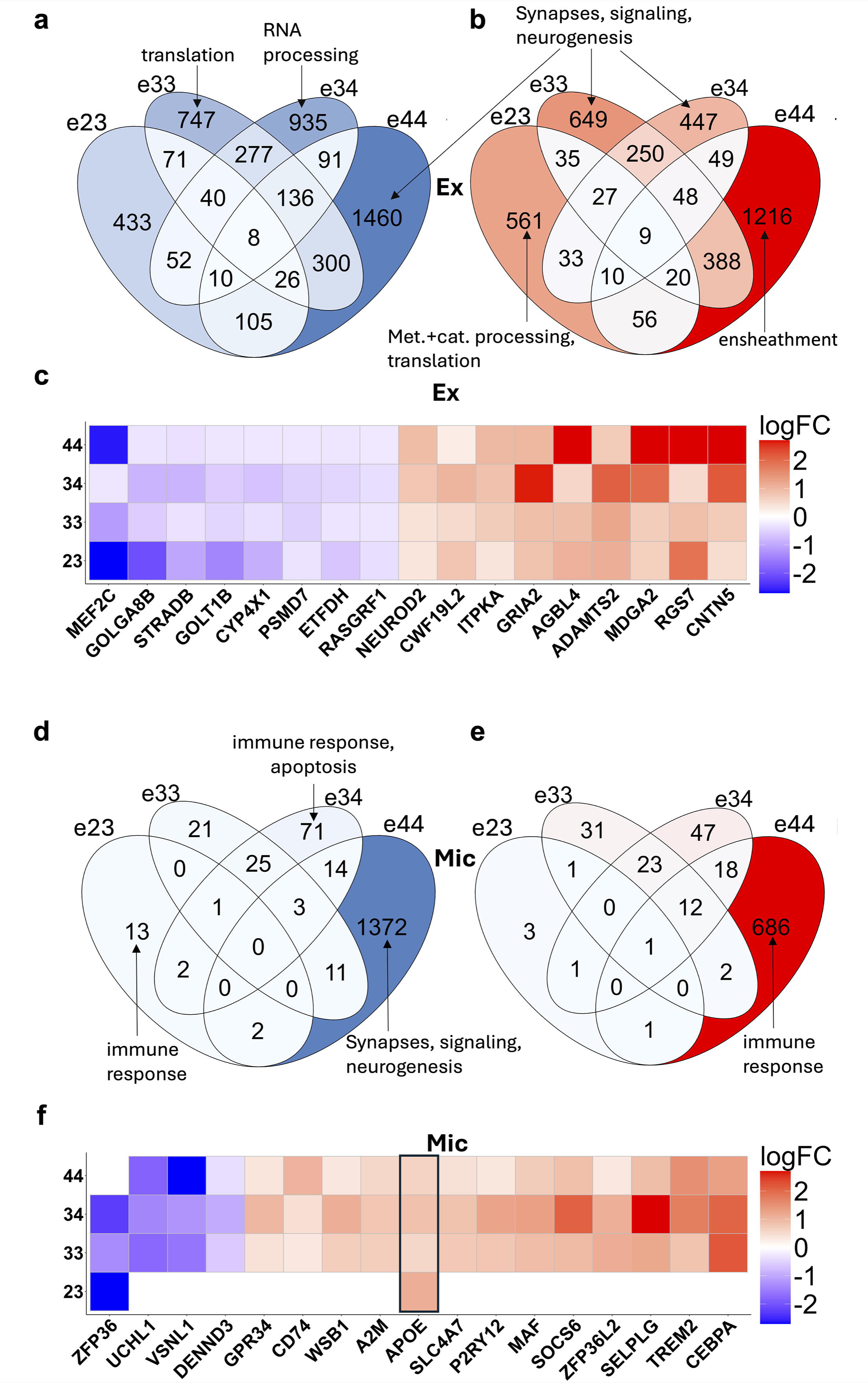
Differentially expressed genes (DEGs) across APOE genotypes in excitatory neurons and microglia. a) Venn diagram showing downregulated DEGs in excitatory neurons, grouped by APOE genotype. Select intersections are annotated with enriched Gene Ontology (GO) terms. b) Venn diagram of upregulated DEGs in excitatory neurons, with enrichment patterns differing by genotype. c) Heatmap of genes commonly upregulated or downregulated across all four genotypes (APOE2/3, APOE3/3, APOE3/4, APOE4/4); color intensity represents log (fold change), bounded between −2 and +2. d) Venn diagram of downregulated DEGs in microglia; e) Venn diagram of upregulated DEGs in microglia; f) Heatmap of DEGs shared across three of the four genotypes in microglia.

Next, we determined genes that are commonly differentially expressed across all four genotypes (*APOE2/3*, *APOE3/3*, *APOE3/4*, and *APOE4/4*) (**Figure 4c**). Eight genes were consistently downregulated (*MEF2C*, *GOLGA8B*, *STRADB*, *CYP4X1*, *PSMD7*, *ETFDH*, and *RASGRF1*), and nine are consistently upregulated (*NEUROD2*, *CWF19L2*, *ITPKA*, *GRIA2*, *AGBL4*, *ADAMTS2*, *MDGA2*, *RGS7*, *CNTN5*) across *APOE* genotypes in excitatory neurons. *MEF2C* has previously been recognized as a gene involved in neuronal vulnerability^126–129^. In **Figure 4c,f**, downregulated genes are displayed in blue and upregulated genes in red, with color intensity indicating the log (fold change). For visualization purposes, values were capped within the range of −2 to +2, with values exceeding these thresholds defaulted to the respective limits. GO analysis of these shared gene sets across APOE genotypes in all major cell types revealed no significantly enriched pathways, regardless of whether the genes were up- or downregulated (**Supplemental Figure 6, 7**). However, the overlaps themselves are statistically significant, with p-values of 2.50 × 10 for the downregulated set and 2.41 × 10 for the upregulated set.

A parallel analysis was conducted for DEGs identified in microglia (**Figure 4d-f**). Overall, the number of DEGs in microglia was considerably lower than in excitatory neurons (**Figure 4d,e**). Like excitatory neurons, *APOE4/4* cases had the highest number of DEGs for both downregulated (**Figure 4d**, 1372) and upregulated (**Figure 4e**, 686) genes. As in excitatory neurons, the genes uniquely downregulated in the *APOE4/4* genotype are enriched for synaptic signaling (*AMPH*, *GABRA1/2/4/5, FLOT1*) and neurogenesis (*OLFM1*, *OLFM3*, *TLR2*) highlighting the importance of glial-neuronal crosstalk in brain function^130^. Additionally, genes downregulated specifically in the *APOE2/3* and *APOE3/4* genotypes are enriched for immune-related processes. In the upregulated gene sets, only the *APOE4/4*-specific genes exhibit significant GO enrichment, which also pertains to immune functions (*C3*, *IL6*, *STAT3*, *TREM2*) (**Supplemental File 4**).

Finally, we conducted an analysis of genes shared across all four genotypes in microglia (**Figure 4f**). The analysis revealed no downregulated genes in common in all four genotypes. A single upregulated gene, *APOE*, was commonly upregulated in all four genotypes. One overlapping gene was shared between three *APOE* genotypes (*APOE2/3*, *APOE3/3*, and *APOE3/4*), *ZFP36* (Zinc Finger Protein 36). All remaining genes in this figure are shared exclusively among *APOE3/3*, *APOE3/4*, and *APOE4/4*.

## DISCUSSION

A wealth of information exists regarding the signaling pathways, transcriptional regulators, and molecular mechanisms involved in AD pathogenesis^131–134^. The recent rise in large scale single cell and spatial transcriptomic data sets has exponentially broadened our view of cell type vulnerability to AD^48^ ^70^ ^135–151^. Recent efforts have focused on single-cell based approaches, with a particular focus on the characterization of transient transcriptional states in relation to AD pathologies^147^ ^152–157^. Here, we have focused on allele variants of the *APOE* gene, the strongest genetic risk factor of AD^158–160^. Our findings demonstrate cell-type specific transcriptional states linked to the presence of the *e4* allele and define AD-associated gene expression changes in all major cell types for each one of the most frequently occurring *APOE* genotypes (*2/3*, *3/3*, *3/4*, and *4/4*).

### APOE’s Role in AD Pathology

It is widely known that *APOE4* is the strongest genetic risk factor for LOAD^34^. A total of 68 unique single-nucleotide polymorphisms (SNPs) have been identified in the human *APOE* sequence^41^; a substantial portion of these SNPs have been directly linked to either increasing or decreasing the risk of developing AD and other neurodegenerative disorders^41^ ^161–165^. Among these, the *e4* allele is the most well-established deleterious variant. Two structural features within the *e4* allelic isoform lead to the protein’s reduced efficacy^3^ ^166^ ^167^. Additionally, a large body of work demonstrates that the *APOE4* protein has toxic gain-of-function effects^168–174^. Efforts to lower *APOE4* expression or function are currently ongoing ^9^ ^175–177^. Accordingly, it is imperative to understand the biological influence of the *APOE* protein and how its allelic variants contribute to AD pathology.

In addition to lipid homeostasis, *APOE* influences inflammation^178–180^, tissue repair^181–183^, and cell viability^174^ ^184–189^. Despite extensive studies, the precise mechanisms by which *APOE* contributes to neurodegeneration remains largely unknown. A long-standing proposed mechanism for how *APOE* contributes to AD vulnerability is through the reduced clearance, increased oligomerization of Aβ, and/or increased amyloid-β precursor protein (APP) transcription ^24^ ^190–197^. Although *APOE* directly binds to Aβ ^160^ ^198–202^, several studies have shown that compared to *APOE2* and *APOE3*, *APOE4* has a lower binding affinity to Aβ ^203–207^. This reduced affinity may permit greater interaction of Aβ with the neuronal plasma membrane, promoting Aβ toxicity and ultimately contributing to neuronal cell damage^208^. In human postmortem studies, more frequent plaques and neurofibrillary tangles are observed in *APOE4* carriers^209^ ^210^. Human PET studies show that the tau accumulation rate is higher in *APOE4* carriers in the parietal, occipital, lateral, and medial temporal cortices^211^ ^212^. However, the mechanisms by which *APOE* contributes to tau pathology appear to be complex. Notably, patients with *APOE4* alleles exhibit heterogeneity in tau accumulation^20^. Murine studies show that *APOE4* drives tau accumulation^6^ ^171^, but may not increase phosphorylated tau pathology^213^. The relationship between *APOE* and AD neuropathology has recently been demonstrated to be more complex by the *APOE3* Christchurch mutation (R136S), where high levels of Aβ, but low levels of tau and resistance to cognitive decline are observed ^163^ ^214^. *APOE3* R136S has been shown to directly bind to tau and suppress its propagation^215^, and to repress microglial response to tau^216^. The mechanism of cognitive resilience in patients harboring *APOE3* R136S has been proposed to be delayed amyloid-facilitated tau pathology^217^.

### e4 Dosage Signatures in Excitatory Neurons are Transcriptionally Distinct from NFT Signatures

*APOE4* disrupts biological mechanisms in a cell-type dependent manner^157^ ^218–220^. In neurons, it has been proposed that *APOE* has direct neurotoxic effects^221^. However, substantial evidence for a broader range of cellular disruptions related to *APOE* has been presented^37^ ^222–226^. Isogenic induced pluripotent stem cell (iPSC)-derived neurons show early neuronal differentiation, increased synapse formation, greater Aβ accumulation and tau phosphorylation in *APOE4* variants compared to *APOE3*^52^.

Excitatory neurons have been considered the most vulnerable cell type to AD pathology^70^ ^227–229^. Excitatory neuronal vulnerability is thought to arise from the preferential accumulation of tau in excitatory versus inhibitory neurons^230–232^, including recently examined across several tauopathies by our group^127^. NFTs, which are intraneuronal aggregates of primarily hyperphosphorylated tau^233–237^, first appear in the dorsal raphe nucleus (DRN) and locus coeruleus (LC) are the first areas to accumulate NFTs^238–242^ and is thought to propagate to the entorhinal cortex and hippocampus in the forebrain followed by association neocortical areas^243–244^. NFT density increases as a function of cognitive decline, as measured by the Clinical Dementia Rating (CDR) scale^245^. Spatiotemporally, in vivo PET studies show that when tau pathology reaches neocortical association areas coincides with the timecourse of onset of significant cognitive decline^243^ ^244^. In addition, some studies have shown a dose-dependent correlation between *APOE4* and NFT burden^30^ ^210^. Given these observations, we examined whether the gene signatures of our excitatory neuronal subclusters overlapped with previously reported NFT-associated signatures. Surprisingly, we found limited shared identity between NFT and *e4*-dosage signatures. Instead, the excitatory neuronal subcluster (ExSub0), which we found to be depleted in *APOE4/4* AD cases, was enriched for NFT signatures. A potential interpretation is that many NFT-bearing neurons have been damaged and/or died due to NFT-related toxicity and the sub-populations that are alive are somewhat resistant neurons with specific signatures. An alternative explanation is that while the presence of NFTs is tightly linked to AD progression ^246–249^, the causative nature of NFTs leading to neuronal cell death has been more recently questioned^235^ ^250^. In primary age-related tauopathy (PART), NFTs deposition is mainly confined to the entorhinal and medial temporal cortex, regions affected early in AD. PART patients typically exhibit normal cognition or mild amnestic changes, with only a minority developing profound impairment that progresses more slowly and less severely than in AD. However, unlike AD, PART is less frequent in *e4*-carriers^251^ ^252^. When examined on a cell-by-cell basis, NFT-bearing neurons do not exhibit typical hallmarks of cell death^55^ ^232^ ^253^ ^254^. Most changes in neuronal gene expression occur prior to NFT formation^254^. Functionally, NFT-positive neurons are responsive to stimuli^255^ ^256^, are morphologically and electrophysiologically comparable^257^, and remain viable ^258^ ^259^. Recently, it has been shown in mouse models that NFT-bearing neurons have increased survival rates compared to non-NFT-bearing neurons^260^. Therefore, NFTs may not be the only pathological factor in neurodegeneration and other species of tau may be more neurotoxic^261–266^. An alternative explanation is that NFT could trigger a very slow process that ultimately leads to neuronal loss. Previous studies directly comparing NFT and non-NFT bearing (“normal”) neurons, demonstrate distinct gene expression patterns^55^ ^267^. NFT-bearing neurons and their immediate surroundings expressed increased levels of proteins involved in autophagy pathways (p62, cathepsin D (CTSD), glucocerebrosidase (GBA), and glycoprotein non-metastatic melanoma protein B (GPNMB)) and vimentin^267^. This agrees with our NFT signature ExSub0 cluster, which also exhibits increased levels of vimentin. Overall, we found limited overlap between *APOE4* dose-dependent and NFT-associated gene signatures, particularly across excitatory neuronal subclusters, suggesting that *APOE4* may drive cognitive decline through mechanisms that are, at least in part, independent of NFT formation. Nevertheless, the enrichment of NFT signatures within the *APOE4*-depleted ExSub0 subcluster reveals a potential area of convergence, indicating that *APOE4* may influence tau-related pathology in specific neuronal contexts.

ExSub0 expresses genes including *GFAP*, *AQP4*, and *S110B*, primarily expressed by astrocytes^268^. However, neuronal *GFAP* expression has been observed in AD patients^269^ and is known to be expressed by neuronal progenitor cells^68^ ^270–272^. Zamboni et al. classified five transcriptional states of astrocytic-derived progenitor cells that follow a distinct pattern of neurogenic trajectory, with early-stage cells expressing genes that significantly overlap with those enriched in our ExSub0 (e.g. *SLC1A3, AQP4, GFAP, NRXN1, NDRG2, CRYAB, CST3, LSAMP*)^67^. The enrichment of neurogenesis genes in *APOE4/4* controls, but not in AD cases, suggests a reprogramming scheme present in a subset of cells in *APOE4/4* controls, that is missing in *APOE4/4* AD patients.

In contrast, ExSub8, depleted in control *e4* carriers, but present in *APOE4/4* AD cases, was enriched for genes involved in synaptic signaling and calcium ion transport. Although there is commonly downregulation of synapse-related genes in vulnerable neuronal subpopulations^70^, enrichment of these genes in ExSub8 was not surprising, as this may be indicative of general disruptions in synaptic homeostasis, which are more prominent in *APOE4/4* AD neurons than in controls. Similarly, enrichment of calcium-related genes in ExSub8 is consistent with existing evidence linking calcium dysfunction to AD^273^ ^274^. Neurons that express high levels of calcium-binding proteins are less vulnerable to AD^275–277^. Calcium homeostasis protects against excitotoxity, a major contributor to excitatory neuronal death^278^. The upregulation of calcium-regulatory genes such as *RYR3*, *CAMK2D/G/N2*, and *MCU* in ExSub8 may reflect compensatory responses to calcium dysregulation in *APOE4/4* AD cases, a response that appears less active or absent in *APOE4/4* controls.

### Microglial Subclustering Revealed Distinct Microglial States

In addition to excitatory neurons, we focused our analyses on microglia. Increased microglial abundance is seen in patients with severe AD pathology^143^. Subpopulations of microglia with distinctive transcriptional signatures, defined as microglial states, have been described in AD^146^ ^147^ ^279^. Previously, it has been difficult to capture microglial activation states related to AD using snRNA-seq analysis from human brain tissue due to depletion of these genes in nuclei compared to whole cells^280^. However, genes that are characteristic to mouse DAM states (*Apoe*, *Trem2*, *Tyrobp*) that are often depleted in microglial nuclei were identified as microglial state markers in this study.

Recent investigations into neurodegenerative microglial states have clarified their role in disease and advanced our overall understanding of glial cell types^281–284^. However, how microglia contribute to disease pathology through *APOE* is not fully understood, despite *APOE* being predominantly secreted by microglia and astrocytes^285–287^. Our microglial subclustering analysis revealed different microglial transcriptional states that differed in proportion based off disease state and/or *APOE* genotype. *HIF1A;APOE*-expressing MicSub8 was upregulated in *e4* carriers. Upregulation of *HIF1A* in microglia is associated with an Aβ plaque-associated microglial state (AβAM)^288^ ^289^ and more generally, is activated under hypoxic conditions including ischemia^290^. Microglial *APOE* upregulation is also associated with Aβ and has been shown to increase tau pathology and neuronal cell death^282^ ^291–293^. Importantly, *APOE* emerged as the only differentially expressed gene (DEG) shared across AD and control samples, regardless of *APOE* genotype, highlighting its central role in microglial function and disease pathogenesis. This falls into alignment with a recent study from Kosoy et al., 2025, which found *APOE* as a top upregulated gene in primary microglia isolated from postmortem AD brains^294^. Another distinct microglial subpopulation, MicSub4 cluster was characterized by high *TREM2* expression. Transition of microglia from a homeostatic to clearance state requires TREM2 activation and BACE1 inhibition^295^. Inhibition of either one of these transitions leads to reduced phagocytic ability ^296,297^. In mouse models, microglial TREM2 deficiency leads to increases in both tau and Aβ pathology, coupled with dysregulated stress kinase signaling in neurons^297^ ^298^. Recently, a new subset of *TREM*-expressing senescent microglia has been defined in 5xFAD mice, which has a distinct transcriptional profile compared to *TREM*-expressing DAM^299^, further emphasizing the heterogeneity and functional specialization of microglial states in AD.

In addition to describing subpopulations of excitatory neurons and microglia across *APOE* genotypes, we also defined a list of *APOE*-dose dependent DEGs. Among these, FRMD4A emerged as a notable candidate, and its expression was further validated using RNAscope. *FRMD4A* (FERM Domain-Containing Protein 4A) was originally found to play a role in epithelial polarization^300^ is a member of the FERM (Four-point-one, Ezrin, Dadixin, Moesin) protein superfamily, whose members have a diverse array of functions^301^. Polymorphisms in *FRMD4A* are associated with AD^302^, and genome-wide association studies have identified it as a risk factor for late-onset Alzheimer’s disease (LOAD)^158^ and cognitive decline in older adults^303^. Furthermore, *FRMD4A* has also been linked to both amyloid and tau release^304–306^. Recently, it has been reported that *FRMD4A* is highly expressed in the microglia of aged human brains^147^. In another recent study, human stem-cell-differentiated microglia (iMGLs) were subject to lentiviral transduction to induce overexpression of transcription factor MITF, a key disease-associated microglia (DAM) regulator and driver of autophagy^293^. RNA-seq analysis of MITF-overexpressing iMGLs showed downregulation of 396 genes, one of which was *FRMD4A*. These findings raise important questions about the potential role of *FRMD4A* in modulating microglial response to neurodegeneration, particularly in the context of *APOE4*. Mechanistic studies to pinpoint the exact role of *FRMD4A* in microglia would be worthwhile.

#### Challenges and Caveats of This Study

A significant challenge when conducting this study is that *APOE* allele frequency is substantially lower for e4, and even more so for e2^39^. The prevalence of certain *APOE* genotype-disease combinations, such as *APOE4/4* control cases is even smaller. Thus, our *APOE4/4* CTRL group was limited to just two cases. Additionally, while our MSSM cohort included *APOE2/2* cases, the neuropathological characteristics of these cases did not fall into defined AD and control groups according to widely accepted criterion. For future studies, it would be interesting to include well-characterized *APOE2/2* and *APOE2/4* cases, as the *e2* allele appears to still confer protection against AD pathology, even in the presence of the *e4* allele^307^.

A caveat of this study is a mild imbalance in sex of the cases included in the integrated cohort (37M, 47F), which can influence transcriptional patterns in both mouse and human ^308–310^. A larger discrepancy is the overwhelming proportion of cases from patients of Caucasian ancestry (12 of 14 cases within our MSSM cohort). According to GWAS, ancestry confers differential risk on e4 vulnerability to AD development ^2^ ^311^. To fully understand the effects of *APOE* allele susceptibility to AD on a global scale, it is essential to increase the representation of individuals from non-European ancestry. As the severity of physiological influence of *APOE* vary across racial groups^312^ and sex^313^, these are important limitations to consider when drawing conclusions about the general population.

Another consideration is that while snRNA-seq is generally a good proxy for transcript abundance of whole cells^314–317^ and is a practical approach for frozen brain tissues^318^ ^319^, some differences between nuclear transcript abundance and whole-cell transcript abundance have been shown. For example, in other tissue types, snRNA-seq has shown lower proportions of immune cells compared with matched scRNA-seq samples^320^. Additionally, droplet-based snRNA-seq may exhibit a bias toward longer transcripts^321^ ^322^. A limitation relating to the RNAscope validation was that, especially for microglia, choosing a microglial-specific marker that is unaltered by disease status is challenging. DAM-microglia downregulate homeostatic markers such as *TMEM119*, *P2RY12* and *SELP1G*^323^. Moreover, it has been recently shown that *TMEM119* is downregulated in microglia in close proximity to amyloid plaques^151^. This may have led to an under-identification of microglia when counting the percent *FRMD4A*+ microglia.

## Conclusion

Currently, the mechanisms through which *APOE4* plays in overall cognitive decline is not fully understood^324–343^. While different isoforms of *APOE* appear to play complex and dynamic roles in cognitive outcomes, it is imperative for us to better understand the underlying cellular mechanisms of *APOE*-driven vulnerability. Here, we comprehensively examined how different *APOE* alleles influence the transcriptional landscape of two highly dynamic and important cell types: excitatory neurons and microglia. We describe subpopulations of these major cell types that are both disease-associated and *APOE*-specific, improving our collective understanding of potential mechanisms underlying susceptibility to AD in patients carrying the *APOE4* allele.

## AWKNOWLEGEMENTS

We would like to thank the Mount Sinai Biorepository and Pathology Core for their assistance with imaging and data analysis. We would also like to thank the brain banks and their funding sources that contributed their resources to this project including MSSM (P30AG066514 ), Emory ADRC grant (P50 AG025688), and UPenn CNDR (P30AG072979, P01AG066597). We are also grateful to the Banner Sun Health Research Institute Brain and Body Donation Program of Sun City, Arizona for the provision of human biological materials. The Brain and Body Donation Program has been supported by the National Institute of Neurological Disorders and Stroke (U24 NS072026 National Brain and Tissue Resource for Parkinson’s Disease and Related Disorders), the National Institute on Aging (P30 AG019610 and P30AG072980, Arizona Alzheimer’s Disease Center), the Arizona Department of Health Services (contract 05700, Arizona Alzheimer’s Research Center), the Arizona Biomedical Research Commission (contracts 4001, 0011, 05-901 and 1001 to the Arizona Parkinson’s Disease Consortium) and the Michael J. Fox Foundation for Parkinson’s Research. We thank Dr. Sivaprakasam Saroja for initial insights in the study.

## AUTHOR CONTRIBUTIONS

Project was designed by ACP, WMW, BZ. Bioinformatics analysis was majorly performed by WMS with contributions from MI, MW, CQC, under BZ supervision. RNAscope was performed by CDS through Mount Sinai Neuropathology Core and analyzed by KM, AC, and RD at Pereira Lab. Post-mortem brains were received from Mount Sinai, UPenn CNDR, Banner Sun Health Institute, and Emory ADRC/CND with further neuropathological insights and contributions from DP and EL. Manuscript was drafted and/or edited by KM, AC, WMS, MI, MW, BZ and ACP. All authors reviewed and approved the manuscript.

## Data and Code Availability

All data supporting the findings of this study are included within the manuscript and its supplementary information files. The scRNA-seq data will be made publicly available at the time of publication. Code used to generate specific figures will be provided upon reasonable request.

## Supplemental Figure Legends

**Supplemental Figure 1.** Reference sample metrics. a) Cluster number score b) Principal component score c) Kolmogorov-Smirnov Score as provided by InPlot() functionality in RISC. The score per sample are shown on the legends on the right. The respective sample annotations are included in Supplemental Table 2. d,e) Identification of RISC PC eigenvectors for d) excitatory neuronal and e) microglial subclustering. d) Variances of RISC eigenvectors within the excitatory neurons. The red line is the mean variances among the 30 RISC eigenvectors. e) X-axis: Eigenvectors identified by RISC for microglial sublustering. Y-axis: The mean values of the eigenvectors across the microglial cells as a proxy for informative features. The red horizontal line marks the cutoff = 2E-5 to distinguish informative features for microglial subclustering.

**Supplemental Figure 2.** Cell type composition and cluster enrichment across three cohorts of samples. a) UMAP plot of cell-type composition for the integrated data set. Legend shows the colors associated with each cohort. b) Cohort-wise enrichment of different cell types in the integrated data set. c) Proportion of cell types enriched for individual samples across all three cohorts. The top legend denotes the genotype and disease status of each sample shown in the graph. The bottom legend shows colors associated with each cell type. d) Fold enrichments of cell types for disease status (AD - left, CTRL – middle, and UNKNOWN - right) for each cohort. The legend denotes the colors associated with each APOE genotype.

**Supplemental Figure 3.** Cohort-wise identification of excitatory neuronal sub clustering. a) Proportion of cells coming from samples from different APOE genotypes by cohort. b) Proportion of excitatory neuronal sub cluster cells from individual cases in three cohorts. Top legend shows color of each excitatory sub cluster shown within bars. Bottom legend shows the color coding for disease status and APOE genotype shown as circles at the top of each bar.

**Supplemental Figure 4.** APOE4/4 P301S mouse cluster 10 is homologous to human NFT-signature Ex5 and ExSub0. a,b) Volcano plots showing module score of each excitatory neuronal cluster enrichment of a) NFT signatures and b) e4 dosage. c) A bar plot shows excitatory-specific NFT signatures for e4 dosage in terms of e44 vs e34, e4 dosage, e4 carrier vs non carrier, and AD vs control. Red line indicates -log10 FET FDR >1.2. d-g) P301S mice harboring APOE4/4 alleles were sacrificed at 9.5 months-of-age and hippocampi were used for snRNA-seq by Wang et al^10^. d) UMAP plot showing major cell populations from APOE4/4 P301S mouse hippocampi. e) Dot plot shows the expression of cell type markers among clusters identified in APOE4/4 P301S mouse hippocampi. f) UMAP plot show overlap between clusters identified in APOE4/4 P301S mouse hippocampi and Ex5 of our integrated human cohort. g) UMAP plot shows NFT signature scores in e4/e4 P301S mouse hippocampi clusters.

**Supplemental Figure 5.** Cohort-wise identification of microglial sub clustering. a) Proportion of cells coming from samples from different APOE genotypes by cohort. b) Proportion of microglial sub cluster cells from individual cases in three cohorts. Top legend shows color of each microglial sub cluster shown within bars. Bottom legend shows the color coding for disease status and APOE genotype shown as circles at the top of each bar.

**Supplemental Figure 6.** Enriched pathways across APOE genotypes in all major cell types. a-g) Venn diagrams showing the number of enriched downregulated (left) and upregulated (right) pathways in all major cell types: a) Ex, b) Mic, c) Ast, d) End, e) In, f) OPC, g) Oli grouped by APOE genotype.

**Supplemental Figure 7.** Enriched DEGs for all major cell types by APOE genotype. a-f) Heatmap of genes commonly upregulated or downregulated across all four genotypes (e23, e33, e34, e44); color intensity represents log (fold change), bounded between −2 and +2.

## Supplemental Files

Supplemental File 1. Extended Neuropath

Supplemental File 2. Integrated Cohort Neuropath Supplemental File 3. Cell type DEGs Supplemental File 4. Cell type GO

Supplemental File 5. Ex_subcluster.markers

Supplemental File 6. MSigDB_results.ExSubcluster_Markers Supplemental File 7. Ex_vs_In.DEG_by_DESeq2 Supplemental File 8. APOE_dosage.signature

Supplemental File 9. Enriched pathways by cluster_e4_dosage_DEG.MSigDB_FET Supplemental File 10. Microglia_subcluster.markers

Supplemental File 11. Microglial Subclustering_GO Supplemental File 12. RNAscope data

## References

1. Belloy ME, Napolioni V, Greicius MD. A Quarter Century of APOE and Alzheimer’s Disease: Progress to Date and the Path Forward. Neuron 2019;101(5):820–38. doi: 10.1016/j.neuron.2019.01.056 [published Online First: 2019/03/08]

2. Farrer LA, Cupples LA, Haines JL, et al. Effects of Age, Sex, and Ethnicity on the Association Between Apolipoprotein E Genotype and Alzheimer Disease: A Meta-analysis. JAMA 1997;278(16):1349–56. doi: 10.1001/jama.1997.03550160069041

3. Mahley RW. Central nervous system lipoproteins: ApoE and regulation of cholesterol metabolism. Arteriosclerosis, thrombosis, and vascular biology 2016;36(7):1305–15.

4. Phillips MC. Apolipoprotein E isoforms and lipoprotein metabolism. IUBMB life 2014;66(9):616–23.

5. Chen Y, Durakoglugil MS, Xian X, et al. ApoE4 reduces glutamate receptor function and synaptic plasticity by selectively impairing ApoE receptor recycling. Proceedings of the National Academy of Sciences 2010;107(26):12011–16.

6. Shi Y, Yamada K, Liddelow SA, et al. ApoE4 markedly exacerbates tau-mediated neurodegeneration in a mouse model of tauopathy. Nature 2017;549(7673):523–27.

7. Shi Y, Manis M, Long J, et al. Microglia drive APOE-dependent neurodegeneration in a tauopathy mouse model. Journal of Experimental Medicine 2019;216(11):2546–61.

8. Shi Y, Andhey PS, Ising C, et al. Overexpressing low-density lipoprotein receptor reduces tau-associated neurodegeneration in relation to apoE-linked mechanisms. Neuron 2021;109(15):2413–26. e7.

9. Litvinchuk A, Huynh TPV, Shi Y, et al. Apolipoprotein E4 reduction with antisense oligonucleotides decreases neurodegeneration in a tauopathy model. Annals of neurology 2021;89(5):952–66.

10. Wang C, Xiong M, Gratuze M, et al. Selective removal of astrocytic APOE4 strongly protects against tau-mediated neurodegeneration and decreases synaptic phagocytosis by microglia. Neuron 2021;109(10):1657–74. e7.

11. Farfel JM, Yu L, De Jager PL, et al. Association of APOE with tau-tangle pathology with and without β-amyloid. Neurobiology of aging 2016;37:19–25.

12. Strittmatter WJ, Weisgraber KH, Huang DY, et al. Binding of human apolipoprotein E to synthetic amyloid beta peptide: isoform-specific effects and implications for late-onset Alzheimer disease. Proceedings of the National Academy of Sciences 1993;90(17):8098–102.

13. Holtzman DM, Bales KR, Tenkova T, et al. Apolipoprotein E isoform-dependent amyloid deposition and neuritic degeneration in a mouse model of Alzheimer’s disease. Proceedings of the National Academy of Sciences 2000;97(6):2892–97. doi: doi:10.1073/pnas.050004797

14. Castellano JM, Kim J, Stewart FR, et al. Human apoE isoforms differentially regulate brain amyloid-β peptide clearance. Science translational medicine 2011;3(89):89ra57-89ra57.

15. Grainger DJ, Reckless J, McKilligin E. Apolipoprotein E modulates clearance of apoptotic bodies in vitro and in vivo, resulting in a systemic proinflammatory state in apolipoprotein E-deficient mice. The Journal of Immunology 2004;173(10):6366–75.

16. Kitagawa K, Matsumoto M, Kuwabara K, et al. Delayed, but marked, expression of apolipoprotein E is involved in tissue clearance after cerebral infarction. Journal of Cerebral Blood Flow & Metabolism 2001;21(10):1199–207.

17. Fagan AM, Murphy BA, Patel SN, et al. Evidence for normal aging of the septo-hippocampal cholinergic system in apoE (−/−) mice but impaired clearance of axonal degeneration products following injury. Experimental neurology 1998;151(2):314–25.

18. Jackson RJ, Hyman BT, Serrano-Pozo A. Multifaceted roles of APOE in Alzheimer disease. Nature Reviews Neurology 2024;20(8):457–74.

19. Jeong W, Lee H, Cho S, et al. ApoE4-Induced Cholesterol Dysregulation and Its Brain Cell Type-Specific Implications in the Pathogenesis of Alzheimer’s Disease. Molecules and Cells 2019;42(11):739–46. doi: 10.14348/molcells.2019.0200

20. Emrani S, Arain HA, DeMarshall C, et al. APOE4 is associated with cognitive and pathological heterogeneity in patients with Alzheimer’s disease: a systematic review. Alzheimer’s Research & Therapy 2020;12(1):141. doi: 10.1186/s13195-020-00712-4

21. Liddell M, Williams J, Bayer A, et al. Confirmation of association between the e4 allele of apolipoprotein E and Alzheimer’s disease. J Med Genet 1994;31(3):197–200. doi: 10.1136/jmg.31.3.197 [published Online First: 1994/03/01]

22. Farlow MR, He Y, Tekin S, et al. Impact of APOE in mild cognitive impairment. Neurology 2004;63(10):1898–901. doi: doi:10.1212/01.WNL.0000144279.21502.B7

23. Reiman EM, Chen K, Alexander GE, et al. Correlations between apolipoprotein E ε4 gene dose and brain-imaging measurements of regional hypometabolism. Proceedings of the National Academy of Sciences 2005;102(23):8299–302. doi: doi:10.1073/pnas.0500579102

24. Huang Y-WA, Zhou B, Wernig M, et al. ApoE2, ApoE3, and ApoE4 differentially stimulate APP transcription and Aβ secretion. Cell 2017;168(3):427–41. e21.

25. Schmechel DE, Saunders AM, Strittmatter WJ, et al. Increased amyloid beta-peptide deposition in cerebral cortex as a consequence of apolipoprotein E genotype in late-onset Alzheimer disease. Proc Natl Acad Sci U S A 1993;90(20):9649–53. doi: 10.1073/pnas.90.20.9649

26. Rebeck GW, Reiter JS, Strickland DK, et al. Apolipoprotein E in sporadic Alzheimer’s disease: allelic variation and receptor interactions. Neuron 1993;11(4):575–80. doi: 10.1016/0896-6273(93)90070-8

27. Fagan AM, Watson M, Parsadanian M, et al. Human and murine ApoE markedly alters A beta metabolism before and after plaque formation in a mouse model of Alzheimer’s disease. Neurobiol Dis 2002;9(3):305–18. doi: 10.1006/nbdi.2002.0483

28. Fouquet M, Besson FL, Gonneaud J, et al. Imaging Brain Effects of APOE4 in Cognitively Normal Individuals Across the Lifespan. Neuropsychology Review 2014;24(3):290–99. doi: 10.1007/s11065-014-9263-8

29. Vanderlip CR, Stark CEL, Initiative tAsDN. APOE4 Increases Susceptibility to Amyloid, Accelerating Episodic Memory Decline. bioRxiv 2024:2024.12.23.630203. doi: 10.1101/2024.12.23.630203

30. Tiraboschi P, Hansen LA, Masliah E, et al. Impact of APOE genotype on neuropathologic and neurochemical markers of Alzheimer disease. Neurology 2004;62(11):1977–83. doi: 10.1212/01.wnl.0000128091.92139.0f

31. Cudaback E, Li X, Montine KS, et al. Apolipoprotein E isoform-dependent microglia migration. Faseb j 2011;25(6):2082–91. doi: 10.1096/fj.10-176891 [published Online First: 20110308]

32. Zhu Y, Nwabuisi-Heath E, Dumanis SB, et al. APOE genotype alters glial activation and loss of synaptic markers in mice. Glia 2012;60(4):559–69. doi: 10.1002/glia.22289 [published Online First: 20120106]

33. Lanfranco MF, Sepulveda J, Kopetsky G, et al. Expression and secretion of apoE isoforms in astrocytes and microglia during inflammation. Glia 2021;69(6):1478–93.

34. Fernández-Calle R, Konings SC, Frontiñán-Rubio J, et al. APOE in the bullseye of neurodegenerative diseases: impact of the APOE genotype in Alzheimer’s disease pathology and brain diseases. Molecular Neurodegeneration 2022;17(1):62. doi: 10.1186/s13024-022-00566-4

35. Tzioras M, Davies C, Newman A, et al. Invited Review: APOE at the interface of inflammation, neurodegeneration and pathological protein spread in Alzheimer’s disease. Neuropathol Appl Neurobiol 2019;45(4):327–46. doi: 10.1111/nan.12529 [published Online First: 20181128]

36. Li Z, Shue F, Zhao N, et al. APOE2: protective mechanism and therapeutic implications for Alzheimer’s disease. Molecular Neurodegeneration 2020;15(1):63. doi: 10.1186/s13024-020-00413-4

37. Guo JL, Braun D, Fitzgerald GA, et al. Decreased lipidated ApoE-receptor interactions confer protection against pathogenicity of ApoE and its lipid cargoes in lysosomes. Cell 2025;188(1):187–206.e26. doi: 10.1016/j.cell.2024.10.027

38. Serrano-Pozo A, Li Z, Noori A, et al. Effect of APOE alleles on the glial transcriptome in normal aging and Alzheimer’s disease. Nat Aging 2021;1(10):919–31. doi: 10.1038/s43587-021-00123-6 [published Online First: 20211011]

39. Lumsden AL, Mulugeta A, Zhou A, et al. Apolipoprotein E (APOE) genotype-associated disease risks: a phenome-wide, registry-based, case-control study utilising the UK Biobank. eBioMedicine 2020;59:102954. doi: 10.1016/j.ebiom.2020.102954

40. Corbo RM, Scacchi R. Apolipoprotein E (APOE) allele distribution in the world. Is APOE* 4 a ‘thrifty’allele? Annals of human genetics 1999;63(4):301–10.

41. Abondio P, Bruno F, Luiselli D. Apolipoprotein E (APOE) haplotypes in healthy subjects from worldwide macroareas: A population genetics perspective for cardiovascular disease, neurodegeneration, and dementia. Current Issues in Molecular Biology 2023;45(4):2817–31.

42. Gao H, Zhang B, Liu L, et al. A universal framework for single-cell multi-omics data integration with graph convolutional networks. Briefings in Bioinformatics 2023;24(3) doi: 10.1093/bib/bbad081

43. Ament SA, Adkins RS, Carter R, et al. The Neuroscience Multi-Omic Archive: a BRAIN Initiative resource for single-cell transcriptomic and epigenomic data from the mammalian brain. Nucleic Acids Research 2022;51(D1):D1075–D85. doi: 10.1093/nar/gkac962

44. Axton M, Baak A, Blomberg N, et al. The FAIR Guiding Principles for scientific data management and stewardship. Scientific data 2016;3(1)

45. de Vries SEJ, Siegle JH, Koch C. Sharing neurophysiology data from the Allen Brain Observatory. eLife 2023;12:e85550. doi: 10.7554/eLife.85550

46. Sunkin SM, Ng L, Lau C, et al. Allen Brain Atlas: an integrated spatio-temporal portal for exploring the central nervous system. Nucleic acids research 2012;41(D1):D996–D1008.

47. Beekly DL, Ramos EM, Lee WW, et al. The National Alzheimer’s Coordinating Center (NACC) database: the uniform data set. Alzheimer Disease & Associated Disorders 2007;21(3):249–58.

48. Mathys H, Davila-Velderrain J, Peng Z, et al. Single-cell transcriptomic analysis of Alzheimer’s disease. Nature 2019;570(7761):332–37. doi: 10.1038/s41586-019-1195-2

49. Zhou Y, Song WM, Andhey PS, et al. Human and mouse single-nucleus transcriptomics reveal TREM2-dependent and TREM2-independent cellular responses in Alzheimer’s disease. Nature Medicine 2020;26(1):131–42. doi: 10.1038/s41591-019-0695-9

50. Forton-Juarez A, Suhy N, Goldman C, et al. Understanding molecular and cellular mechanisms of APOE4-mediated susceptibility to tau-related cognitive impairments in the miBrain model. Alzheimer’s & Dementia 2023;19(S13):e080238. doi: 10.1002/alz.080238

51. Haney MS, Pálovics R, Munson CN, et al. APOE4/4 is linked to damaging lipid droplets in Alzheimer’s disease microglia. Nature 2024;628(8006):154–61. doi: 10.1038/s41586-024-07185-7

52. Lin Y-T, Seo J, Gao F, et al. APOE4 Causes Widespread Molecular and Cellular Alterations Associated with Alzheimer’s Disease Phenotypes in Human iPSC-Derived Brain Cell Types. Neuron 2018;98(6):1141–54.e7. doi: 10.1016/j.neuron.2018.05.008

53. Preman P, Moechars D, Fertan E, et al. APOE from astrocytes restores Alzheimer’s Aβ-pathology and DAM-like responses in APOE deficient microglia. EMBO Molecular Medicine 2024;16(12):3113–41. doi: 10.1038/s44321-024-00162-7

54. Rao A, Chen N, Kim MJ, et al. Microglia depletion reduces human neuronal APOE4-related pathologies in a chimeric Alzheimer&#x2019;s disease model. Cell Stem Cell 2025;32(1):86–104.e7. doi: 10.1016/j.stem.2024.10.005

55. Otero-Garcia M, Mahajani SU, Wakhloo D, et al. Molecular signatures underlying neurofibrillary tangle susceptibility in Alzheimer’s disease. Neuron 2022;110(18):2929–48.e8. doi: 10.1016/j.neuron.2022.06.021 [published Online First: 20220725]

56. Beach TG, Adler CH, Sue LI, et al. Arizona Study of Aging and Neurodegenerative Disorders and Brain and Body Donation Program. Neuropathology 2015;35(4):354–89. doi: 10.1111/neup.12189 [published Online First: 20150126]

57. Montine TJ, Phelps CH, Beach TG, et al. National Institute on Aging-Alzheimer’s Association guidelines for the neuropathologic assessment of Alzheimer’s disease: a practical approach. Acta Neuropathol 2012;123(1):1–11. doi: 10.1007/s00401-011-0910-3 [published Online First: 20111120]

58. Stuart T, Butler A, Hoffman P, et al. Comprehensive integration of single-cell data. Cell 2019;177(7):1888–902. e21.

59. Finak G, McDavid A, Yajima M, et al. MAST: a flexible statistical framework for assessing transcriptional changes and characterizing heterogeneity in single-cell RNA sequencing data. Genome biology 2015;16(1):1–13.

60. McGinnis CS, Murrow LM, Gartner ZJ. DoubletFinder: doublet detection in single-cell RNA sequencing data using artificial nearest neighbors. Cell systems 2019;8(4):329–37. e4.

61. Liu Y, Wang T, Zhou B, et al. Robust integration of multiple single-cell RNA sequencing datasets using a single reference space. Nature biotechnology 2021;39(7):877–84.

62. Love MI, Huber W, Anders S. Moderated estimation of fold change and dispersion for RNA-seq data with DESeq2. Genome Biology 2014;15(12):550. doi: 10.1186/s13059-014-0550-8

63. McGinnis CS, Murrow LM, Gartner ZJ. DoubletFinder: Doublet Detection in Single-Cell RNA Sequencing Data Using Artificial Nearest Neighbors. Cell Syst 2019;8(4):329–37 e4. doi: 10.1016/j.cels.2019.03.003 [published Online First: 2019/04/08]

64. Stuart T, Butler A, Hoffman P, et al. Comprehensive Integration of Single-Cell Data. Cell 2019;177(7):1888–902 e21. doi: 10.1016/j.cell.2019.05.031 [published Online First: 2019/06/11]

65. Finak G, McDavid A, Yajima M, et al. MAST: a flexible statistical framework for assessing transcriptional changes and characterizing heterogeneity in single-cell RNA sequencing data. Genome Biol 2015;16:278. doi: 10.1186/s13059-015-0844-5 [published Online First: 2015/12/15]

66. Subramanian A, Tamayo P, Mootha VK, et al. Gene set enrichment analysis: a knowledge-based approach for interpreting genome-wide expression profiles. Proc Natl Acad Sci U S A 2005;102(43):15545–50. doi: 10.1073/pnas.0506580102 [published Online First: 2005/10/04]

67. Zamboni M, Llorens-Bobadilla E, Magnusson JP, et al. A Widespread Neurogenic Potential of Neocortical Astrocytes Is Induced by Injury. Cell Stem Cell 2020;27(4):605–17.e5. doi: 10.1016/j.stem.2020.07.006

68. Garcia ADR, Doan NB, Imura T, et al. GFAP-expressing progenitors are the principal source of constitutive neurogenesis in adult mouse forebrain. Nature Neuroscience 2004;7(11):1233–41. doi: 10.1038/nn1340

69. Wu Y, Korobeynyk VI, Zamboni M, et al. Multimodal transcriptomics reveal neurogenic aging trajectories and age-related regional inflammation in the dentate gyrus. Nature Neuroscience 2025 doi: 10.1038/s41593-024-01848-4

70. Leng K, Li E, Eser R, et al. Molecular characterization of selectively vulnerable neurons in Alzheimer’s disease. Nat Neurosci 2021;24(2):276–87. doi: 10.1038/s41593-020-00764-7 [published Online First: 2021/01/13]

71. Nakaya N, Sultana A, Lee H-S, et al. Olfactomedin 1 interacts with the Nogo A receptor complex to regulate axon growth. Journal of Biological Chemistry 2012;287(44):37171–84.

72. Nakaya N, Lee H-S, Takada Y, et al. Zebrafish olfactomedin 1 regulates retinal axon elongation in vivo and is a modulator of Wnt signaling pathway. Journal of Neuroscience 2008;28(31):7900–10.

73. Nakaya N, Sultana A, Munasinghe J, et al. Deletion in the N-terminal half of olfactomedin 1 modifies its interaction with synaptic proteins and causes brain dystrophy and abnormal behavior in mice. Experimental neurology 2013;250:205–18.

74. Manganas LN, Durá I, Osenberg S, et al. BASP1 labels neural stem cells in the neurogenic niches of mammalian brain. Scientific Reports 2021;11(1):5546. doi: 10.1038/s41598-021-85129-1

75. Caroni P, Aigner L, Schneider C. Intrinsic neuronal determinants locally regulate extrasynaptic and synaptic growth at the adult neuromuscular junction. The Journal of cell biology 1997;136(3):679–92.

76. Hartl M, Schneider R. A Unique Family of Neuronal Signaling Proteins Implicated in Oncogenesis and Tumor Suppression. Front Oncol 2019;9:289. doi: 10.3389/fonc.2019.00289 [published Online First: 20190417]

77. Cool DR, Normant E, Shen F-s, et al. Carboxypeptidase E is a regulated secretory pathway sorting receptor: genetic obliteration leads to endocrine disorders in Cpefat mice. Cell 1997;88(1):73–83.

78. Lou H, Kim S-K, Zaitsev E, et al. Sorting and activity-dependent secretion of BDNF require interaction of a specific motif with the sorting receptor carboxypeptidase e. Neuron 2005;45(2):245–55.

79. Lyons CE, Zhou X, Razzoli M, et al. Lifelong chronic psychosocial stress induces a proteomic signature of Alzheimer’s disease in wildtype mice. European Journal of Neuroscience 2022;55(9-10):2971–85. doi: 10.1111/ejn.15329

80. Barranco N, Plá V, Alcolea D, et al. Dense core vesicle markers in CSF and cortical tissues of patients with Alzheimer’s disease. Transl Neurodegener 2021;10(1):37. doi: 10.1186/s40035-021-00263-0 [published Online First: 20210926]

81. Plá V, Paco S, Ghezali G, et al. Secretory Sorting Receptors Carboxypeptidase E and Secretogranin III in Amyloid β-Associated Neural Degeneration in Alzheimer’s Disease. Brain Pathology 2013;23(3):274–84. doi: 10.1111/j.1750-3639.2012.00644.x

82. Xiao L, Loh YP. Neurotrophic Factor-α1/Carboxypeptidase E Functions in Neuroprotection and Alleviates Depression. Front Mol Neurosci 2022;15:918852. doi: 10.3389/fnmol.2022.918852 [published Online First: 20220526]

83. Xiao L, Yang X, Loh YP. Neurotrophic, Gene Regulation, and Cognitive Functions of Carboxypeptidase E-Neurotrophic Factor-α1 and Its Variants. Front Neurosci 2019;13:243. doi: 10.3389/fnins.2019.00243 [published Online First: 20190319]

84. Cheng Y, Cawley NX, Loh YP. Carboxypeptidase E/NFα1: a new neurotrophic factor against oxidative stress-induced apoptotic cell death mediated by ERK and PI3-K/AKT pathways. PloS one 2013;8(8):e71578.

85. Fan F-C, Du Y, Zheng W-H, et al. Carboxypeptidase E conditional knockout mice exhibit learning and memory deficits and neurodegeneration. Translational Psychiatry 2023;13(1):135. doi: 10.1038/s41398-023-02429-y

86. Zhao K, Zhang M, Zhang L, et al. Intracellular osteopontin stabilizes TRAF3 to positively regulate innate antiviral response. Scientific Reports 2016;6(1):23771. doi: 10.1038/srep23771

87. Liu E, Sun J, Yang J, et al. ZDHHC11 positively regulates NF-κB activation by enhancing TRAF6 oligomerization. Frontiers in Cell and Developmental Biology 2021;9:710967.

88. Liu Y, Zhou Q, Zhong L, et al. ZDHHC11 modulates innate immune response to DNA virus by mediating MITA–IRF3 association. Cellular & molecular immunology 2018;15(10):907–16.

89. Du J, Luo H, Ye S, et al. Unraveling IFI44L’s biofunction in human disease. Frontiers in Oncology 2024;Volume 14 - 2024 doi: 10.3389/fonc.2024.1436576

90. Volkert MR, Elliott NA, Housman DE. Functional genomics reveals a family of eukaryotic oxidation protection genes. Proc Natl Acad Sci U S A 2000;97(26):14530–5. doi: 10.1073/pnas.260495897

91. Elliott NA, Volkert MR. Stress induction and mitochondrial localization of Oxr1 proteins in yeast and humans. Mol Cell Biol 2004;24(8):3180–7. doi: 10.1128/mcb.24.8.3180-3187.2004

92. Andrade WA, Silva AM, Alves VS, et al. Early endosome localization and activity of RasGEF1b, a toll-like receptor-inducible Ras guanine-nucleotide exchange factor. Genes & Immunity 2010;11(6):447–57. doi: 10.1038/gene.2009.107

93. Woo SH, Lukacs V, de Nooij JC, et al. Piezo2 is the principal mechanotransduction channel for proprioception. Nat Neurosci 2015;18(12):1756–62. doi: 10.1038/nn.4162 [published Online First: 20151109]

94. Schroeder BC, Hechenberger M, Weinreich F, et al. KCNQ5, a Novel Potassium Channel Broadly Expressed in Brain, Mediates M-type Currents*. Journal of Biological Chemistry 2000;275(31):24089–95. doi: 10.1074/jbc.M003245200

95. Sturchler E, Cox JA, Durussel I, et al. S100A16, a novel calcium-binding protein of the EF-hand superfamily. J Biol Chem 2006;281(50):38905–17. doi: 10.1074/jbc.M605798200 [published Online First: 20061008]

96. Piirsoo M, Meijer D, Timmusk T. Expression analysis of the CLCA gene family in mouse and human with emphasis on the nervous system. BMC Developmental Biology 2009;9(1):10. doi: 10.1186/1471-213X-9-10

97. Lalonde R, Strazielle C. The DST gene in neurobiology. Journal of Neurogenetics 2023;37(4):131–38. doi: 10.1080/01677063.2024.2319880

98. Csukasi F, Duran I, Barad M, et al. The PTH/PTHrP-SIK3 pathway affects skeletogenesis through altered mTOR signaling. Sci Transl Med 2018;10(459) doi: 10.1126/scitranslmed.aat9356

99. Shahidin, Wang Y, Wu Y, et al. Selenium and Selenoproteins: Mechanisms, Health Functions, and Emerging Applications. Molecules 2025;30(3):437.

100. Li Y, Zhou Y, Liu D, et al. Glutathione Peroxidase 3 induced mitochondria-mediated apoptosis via AMPK/ERK1/2 pathway and resisted autophagy-related ferroptosis via AMPK/mTOR pathway in hyperplastic prostate. Journal of Translational Medicine 2023;21(1):575.

101. Abner EL, Neltner JH, Jicha GA, et al. Diffuse Amyloid-β Plaques, Neurofibrillary Tangles, and the Impact of APOE in Elderly Persons’ Brains Lacking Neuritic Amyloid Plaques. J Alzheimers Dis 2018;64(4):1307–24. doi: 10.3233/jad-180514

102. Sparks DL, Scheff SW, Liu H, et al. Increased density of senile plaques (SP), but not neurofibrillary tangles (NFT), in non-demented individuals with the apolipoprotein E4 allele: comparison to confirmed Alzheimer’s disease patients. Journal of the Neurological Sciences 1996;138(1):97–104. doi: 10.1016/0022-510X(96)00008-1

103. Budny V, Ruminot I, Wybitul M, et al. Fueling the brain - the role of apolipoprotein E in brain energy metabolism and its implications for Alzheimer’s disease. Translational Psychiatry 2025;15(1):316. doi: 10.1038/s41398-025-03550-w

104. Keren-Shaul H, Spinrad A, Weiner A, et al. A Unique Microglia Type Associated with Restricting Development of Alzheimer’s Disease. Cell 2017;169(7):1276–90.e17. doi: 10.1016/j.cell.2017.05.018 [published Online First: 20170608]

105. Millet A, Ledo JH, Tavazoie SF. An exhausted-like microglial population accumulates in aged and APOE4 genotype Alzheimer’s brains. Immunity 2024;57(1):153–70.e6. doi: 10.1016/j.immuni.2023.12.001

106. Tang Y, Hu H, Xie Q, et al. GAS6/AXL signaling promotes M2 microglia efferocytosis to alleviate neuroinflammation in sepsis-associated encephalopathy. Cell Death Discovery 2025;11(1):268. doi: 10.1038/s41420-025-02507-8

107. Dongre P, Ramesh M, Govindaraju T, et al. Asrij/OCIAD1 depletion reduces inflammatory microglial activation and ameliorates Aβ pathology in an Alzheimer’s disease mouse model. Journal of Neuroinflammation 2025;22(1):89. doi: 10.1186/s12974-025-03415-5

108. Kim S-M, Mun B-R, Lee S-J, et al. TREM2 promotes Aβ phagocytosis by upregulating C/EBPα-dependent CD36 expression in microglia. Scientific Reports 2017;7(1):11118. doi: 10.1038/s41598-017-11634-x

109. Akhter R, Shao Y, Formica S, et al. TREM2 alters the phagocytic, apoptotic and inflammatory response to Aβ(42) in HMC3 cells. Mol Immunol 2021;131:171–79. doi: 10.1016/j.molimm.2020.12.035 [published Online First: 20210115]

110. Hwang M, Savarin C, Kim J, et al. Trem2 deficiency impairs recovery and phagocytosis and dysregulates myeloid gene expression during virus-induced demyelination. Journal of Neuroinflammation 2022;19(1):267. doi: 10.1186/s12974-022-02629-1

111. Meng J, Han L, Xu H, et al. TREM2 regulates microglial phagocytosis of synapses in innate immune tolerance. Int Immunopharmacol 2024;127:111445. doi: 10.1016/j.intimp.2023.111445 [published Online First: 20231225]

112. Damisah EC, Rai A, Grutzendler J. TREM2: Modulator of Lipid Metabolism in Microglia. Neuron 2020;105(5):759–61. doi: 10.1016/j.neuron.2020.02.008

113. Nugent AA, Lin K, van Lengerich B, et al. TREM2 Regulates Microglial Cholesterol Metabolism upon Chronic Phagocytic Challenge. Neuron 2020;105(5):837–54.e9. doi: 10.1016/j.neuron.2019.12.007

114. Wang Y, Cella M, Mallinson K, et al. TREM2 lipid sensing sustains the microglial response in an Alzheimer’s disease model. Cell 2015;160(6):1061–71.

115. Sun N, Victor MB, Park YP, et al. Human microglial state dynamics in Alzheimer’s disease progression. Cell 2023;186(20):4386–403 e29. doi: 10.1016/j.cell.2023.08.037 [published Online First: 2023/09/30]

116. Lee D, Porras C, Spencer C, et al. Plasticity of Human Microglia and Brain Perivascular Macrophages in Aging and Alzheimer’s Disease. medRxiv 2023:2023.10.25.23297558. doi: 10.1101/2023.10.25.23297558

117. Saito K, Mori M, Kambara N, et al. FilGAP, a GAP protein for Rac, regulates front rear polarity and tumor cell migration through the ECM. The FASEB Journal 2021;35(4):e21508.

118. Lavelin I, Geiger B. Characterization of a Novel GTPase-activating Protein Associated with Focal Adhesions and the Actin Cytoskeleton*. Journal of Biological Chemistry 2005;280(8):7178–85. doi: 10.1074/jbc.M411990200

119. Mori M, Saito K, Ohta Y. ARHGAP22 localizes at endosomes and regulates actin cytoskeleton. PLoS One 2014;9(6):e100271.

120. Longatti A, Ponzoni L, Moretto E, et al. Arhgap22 disruption leads to RAC1 hyperactivity affecting hippocampal glutamatergic synapses and cognition in mice. Molecular Neurobiology 2021;58(12):6092–110.

121. Seoh ML, Ng CH, Yong J, et al. ArhGAP15, a novel human RacGAP protein with GTPase binding property. FEBS Lett 2003;539(1-3):131–7. doi: 10.1016/s0014-5793(03)00213-8

122. Sun Z, Zhang B, Wang C, et al. Forkhead box P3 regulates ARHGAP 15 expression and affects migration of glioma cells through the Rac1 signaling pathway. Cancer science 2017;108(1):61–72.

123. Yang W, Wang B, Yu Q, et al. ARHGAP24 represses β-catenin transactivation-induced invasiveness in hepatocellular carcinoma mainly by acting as a GTPase-independent scaffold. Theranostics 2022;12(14):6189.

124. McPhie DL, Coopersmith R, Hines-Peralta A, et al. DNA synthesis and neuronal apoptosis caused by familial Alzheimer disease mutants of the amyloid precursor protein are mediated by the p21 activated kinase PAK3. J Neurosci 2003;23(17):6914–27. doi: 10.1523/jneurosci.23-17-06914.2003

125. Ge Y, Zhou L, Fu Y, et al. Caspase-2 is a condensate-mediated deubiquitinase in protein quality control. Nature Cell Biology 2024;26(11):1943–57. doi: 10.1038/s41556-024-01522-8

126. Castanho I, Yeganeh PN, Boix CA, et al. Molecular hallmarks of excitatory and inhibitory neuronal resilience and resistance to Alzheimer’s disease. bioRxiv 2025:2025.01.13.632801. doi: 10.1101/2025.01.13.632801

127. Broekaart DWM, Sharma A, Ramakrishnan A, et al. Molecular signatures of regional vulnerability to tauopathy in excitatory cortical neurons. Acta Neuropathol 2025;149(1):60. doi: 10.1007/s00401-025-02879-2 [published Online First: 20250607]

128. Udeochu JC, Amin S, Huang Y, et al. Tau activation of microglial cGAS–IFN reduces MEF2C-mediated cognitive resilience. Nature Neuroscience 2023;26(5):737–50. doi: 10.1038/s41593-023-01315-6

129. Barker SJ, Raju RM, Milman NEP, et al. MEF2 is a key regulator of cognitive potential and confers resilience to neurodegeneration. Science Translational Medicine 2021;13(618):eabd7695. doi: doi:10.1126/scitranslmed.abd7695

130. Marinelli S, Basilico B, Marrone MC, et al. Microglia-neuron crosstalk: Signaling mechanism and control of synaptic transmission. Semin Cell Dev Biol 2019;94:138–51. doi: 10.1016/j.semcdb.2019.05.017 [published Online First: 20190530]

131. Godoy JA, Rios JA, Zolezzi JM, et al. Signaling pathway cross talk in Alzheimer’s disease. Cell Communication and Signaling 2014;12(1):23. doi: 10.1186/1478-811X-12-23

132. Mizuno S, Iijima R, Ogishima S, et al. AlzPathway: a comprehensive map of signaling pathways of Alzheimer’s disease. BMC Systems Biology 2012;6(1):52. doi: 10.1186/1752-0509-6-52

133. Ho GJ, Drego R, Hakimian E, et al. Mechanisms of cell signaling and inflammation in Alzheimer’s disease. Current Drug Targets-Inflammation & Allergy 2005;4(2):247–56.

134. Gadhave K, Kumar D, Uversky VN, et al. A multitude of signaling pathways associated with Alzheimer’s disease and their roles in AD pathogenesis and therapy. Medicinal research reviews 2021;41(5):2689–745.

135. Bagyinszky E, Giau VV, An SA. Transcriptomics in Alzheimer’s Disease: Aspects and Challenges. Int J Mol Sci 2020;21(10) doi: 10.3390/ijms21103517 [published Online First: 20200515]

136. Cain A, Taga M, McCabe C, et al. Multicellular communities are perturbed in the aging human brain and Alzheimer’s disease. Nature neuroscience 2023;26(7):1267–80.

137. Gabitto MI, Travaglini KJ, Rachleff VM, et al. Integrated multimodal cell atlas of Alzheimer’s disease. Nature Neuroscience 2024;27(12):2366–83. doi: 10.1038/s41593-024-01774-5

138. Green GS, Yang HS, Fujita M, et al. Cellular dynamics across aged human brains uncover a multicellular cascade leading to Alzheimer’s disease. Alzheimer’s & Dementia 2023;19:e083212.

139. Green GS, Fujita M, Yang H-S, et al. Cellular communities reveal trajectories of brain ageing and Alzheimer’s disease. Nature 2024;633(8030):634–45. doi: 10.1038/s41586-024-07871-6

140. Grubman A, Chew G, Ouyang JF, et al. A single-cell atlas of entorhinal cortex from individuals with Alzheimer’s disease reveals cell-type-specific gene expression regulation. Nature Neuroscience 2019;22(12):2087–97. doi: 10.1038/s41593-019-0539-4

141. Lau S-F, Cao H, Fu AK, et al. Single-nucleus transcriptome analysis reveals dysregulation of angiogenic endothelial cells and neuroprotective glia in Alzheimer’s disease. Proceedings of the National Academy of Sciences 2020;117(41):25800–09.

142. Liu A, Fernandes BS, Citu C, et al. Unraveling the intercellular communication disruption and key pathways in Alzheimer’s disease: an integrative study of single-nucleus transcriptomes and genetic association. Alzheimer’s Research & Therapy 2024;16(1):3. doi: 10.1186/s13195-023-01372-w

143. Mathys H, Peng Z, Boix CA, et al. Single-cell atlas reveals correlates of high cognitive function, dementia, and resilience to Alzheimer&#x2019;s disease pathology. Cell 2023;186(20):4365–85.e27. doi: 10.1016/j.cell.2023.08.039

144. Mathys H, Boix CA, Akay LA, et al. Single-cell multiregion dissection of Alzheimer’s disease. Nature 2024;632(8026):858–68. doi: 10.1038/s41586-024-07606-7

145. Morabito S, Miyoshi E, Michael N, et al. Single-nucleus chromatin accessibility and transcriptomic characterization of Alzheimer’s disease. Nature genetics 2021;53(8):1143–55.

146. Prater KE, Green KJ, Mamde S, et al. Human microglia show unique transcriptional changes in Alzheimer’s disease. Nature Aging 2023;3(7):894–907.

147. Sun N, Victor MB, Park YP, et al. Human microglial state dynamics in Alzheimer’s disease progression. Cell 2023;186(20):4386–403.e29. doi: 10.1016/j.cell.2023.08.037

148. Wang Q, Antone J, Alsop E, et al. Single cell transcriptomes and multiscale networks from persons with and without Alzheimer’s disease. Nat Commun 2024;15(1):5815. doi: 10.1038/s41467-024-49790-0 [published Online First: 20240710]

149. Dharshini SAP, Sanz-Ros J, Pan J, et al. Molecular Signatures of Resilience to Alzheimer’s Disease in Neocortical Layer 4 Neurons. bioRxiv 2024:2024.11.03.621787. doi: 10.1101/2024.11.03.621787

150. Huuki-Myers LA, Spangler A, Eagles NJ, et al. A data-driven single-cell and spatial transcriptomic map of the human prefrontal cortex. Science 2024;384(6698):eadh1938. doi: doi:10.1126/science.adh1938

151. Mallach A, Zielonka M, van Lieshout V, et al. Microglia-astrocyte crosstalk in the amyloid plaque niche of an Alzheimer&#x2019;s disease mouse model, as revealed by spatial transcriptomics. Cell Reports 2024;43(6) doi: 10.1016/j.celrep.2024.114216

152. Olah M, Menon V, Habib N, et al. Single cell RNA sequencing of human microglia uncovers a subset associated with Alzheimer’s disease. Nature Communications 2020;11(1):6129. doi: 10.1038/s41467-020-19737-2

153. Smith AM, Davey K, Tsartsalis S, et al. Diverse human astrocyte and microglial transcriptional responses to Alzheimer’s pathology. Acta Neuropathol 2022;143(1):75–91. doi: 10.1007/s00401-021-02372-6 [published Online First: 20211112]

154. Gerrits E, Brouwer N, Kooistra SM, et al. Distinct amyloid-β and tau-associated microglia profiles in Alzheimer’s disease. Acta Neuropathol 2021;141(5):681–96. doi: 10.1007/s00401-021-02263-w [published Online First: 20210220]

155. Sayed FA, Kodama L, Fan L, et al. AD-linked R47H-TREM2 mutation induces disease-enhancing microglial states via AKT hyperactivation. Sci Transl Med 2021;13(622):eabe3947. doi: 10.1126/scitranslmed.abe3947 [published Online First: 20211201]

156. Gazestani V, Kamath T, Nadaf NM, et al. Early Alzheimer’s disease pathology in human cortex involves transient cell states. Cell 2023;186(20):4438–53.e23. doi: 10.1016/j.cell.2023.08.005

157. Blanchard JW, Akay LA, Davila-Velderrain J, et al. APOE4 impairs myelination via cholesterol dysregulation in oligodendrocytes. Nature 2022;611(7937):769–79. doi: 10.1038/s41586-022-05439-w

158. Lambert JC, Ibrahim-Verbaas CA, Harold D, et al. Meta-analysis of 74,046 individuals identifies 11 new susceptibility loci for Alzheimer’s disease. Nat Genet 2013;45(12):1452–8. doi: 10.1038/ng.2802 [published Online First: 20131027]

159. Corder EH, Saunders AM, Strittmatter WJ, et al. Gene dose of apolipoprotein E type 4 allele and the risk of Alzheimer’s disease in late onset families. Science 1993;261(5123):921–23.

160. Strittmatter WJ, Saunders AM, Schmechel D, et al. Apolipoprotein E: high-avidity binding to beta-amyloid and increased frequency of type 4 allele in late-onset familial Alzheimer disease. Proceedings of the National Academy of Sciences 1993;90(5):1977–81.

161. Seripa D, D’Onofrio G, Panza F, et al. The genetics of the human APOE polymorphism. Rejuvenation research 2011;14(5):491–500.

162. Namboori PK, Vineeth K, Rohith V, et al. The ApoE gene of Alzheimer’s disease (AD). Functional & integrative genomics 2011;11:519–22.

163. Arboleda-Velasquez JF, Lopera F, O’Hare M, et al. Resistance to autosomal dominant Alzheimer’s disease in an APOE3 Christchurch homozygote: a case report. Nature Medicine 2019;25(11):1680–83. doi: 10.1038/s41591-019-0611-3

164. Almeida MC, Eger SJ, He C, et al. Single-nucleus RNA sequencing demonstrates an autosomal dominant Alzheimer’s disease profile and possible mechanisms of disease protection. Neuron 2024;112(11):1778–94.e7. doi: 10.1016/j.neuron.2024.02.009

165. Gharbi-Meliani A, Dugravot A, Sabia S, et al. The association of APOE ε4 with cognitive function over the adult life course and incidence of dementia: 20 years follow-up of the Whitehall II study. Alzheimer’s research & therapy 2021;13:1–11.

166. Hatters DM, Peters-Libeu CA, Weisgraber KH. Apolipoprotein E structure: insights into function. Trends in Biochemical Sciences 2006;31(8):445–54. doi: 10.1016/j.tibs.2006.06.008

167. Mahley RW, Huang Y. Apolipoprotein (apo) E4 and Alzheimer’s disease: unique conformational and biophysical properties of apoE4 can modulate neuropathology. Acta Neurologica Scandinavica 2006;114:8–14.

168. Rieker C, Migliavacca E, Vaucher A, et al. Apolipoprotein E4 Expression Causes Gain of Toxic Function in Isogenic Human Induced Pluripotent Stem Cell-Derived Endothelial Cells. *Arteriosclerosis*, Thrombosis, and Vascular Biology 2019;39(9):e195–e207. doi: doi:10.1161/ATVBAHA.118.312261

169. Wang C, Najm R, Xu Q, et al. Gain of toxic apolipoprotein E4 effects in human iPSC-derived neurons is ameliorated by a small-molecule structure corrector. Nat Med 2018;24(5):647–57. doi: 10.1038/s41591-018-0004-z [published Online First: 20180409]

170. Zepa L, Frenkel M, Belinson H, et al. ApoE4 Driven Accumulation of Intraneuronal Oligomerized Aβ42 following Activation of the Amyloid Cascade In Vivo Is Mediated by a Gain of Function. International Journal of Alzheimer’s Disease 2011;2011(1):792070.

171. Liraz O, Boehm-Cagan A, Michaelson DM. ApoE4 induces Aβ42, tau, and neuronal pathology in the hippocampus of young targeted replacement apoE4 mice. Molecular neurodegeneration 2013;8:1–17.

172. Zhao N, Attrebi ON, Ren Y, et al. APOE4 exacerbates α-synuclein pathology and related toxicity independent of amyloid. Science translational medicine 2020;12(529):eaay1809.

173. Najm R, Zalocusky KA, Zilberter M, et al. In vivo chimeric Alzheimer’s disease modeling of apolipoprotein E4 toxicity in human neurons. Cell reports 2020;32(4)

174. Wadhwani AR, Affaneh A, Van Gulden S, et al. Neuronal apolipoprotein E4 increases cell death and phosphorylated tau release in alzheimer disease. Annals of Neurology 2019;85(5):726–39. doi: 10.1002/ana.25455

175. Vance JM, Farrer LA, Huang Y, et al. Report of the APOE4 National Institute on Aging/Alzheimer Disease Sequencing Project Consortium Working Group: Reducing APOE4 in Carriers is a Therapeutic Goal for Alzheimer’s Disease. Annals of Neurology 2024;95&lt;otherinfo>(4)&lt;/otherinfo>:625-34. doi: 10.1002/ana.26864

176. Uddin MS, Kabir MT, Al Mamun A, et al. APOE and Alzheimer’s disease: evidence mounts that targeting APOE4 may combat Alzheimer’s pathogenesis. Molecular neurobiology 2019;56(4):2450–65.

177. Li Y, Macyczko JR, Liu C-C, et al. ApoE4 reduction: An emerging and promising therapeutic strategy for Alzheimer’s disease. Neurobiology of aging 2022;115:20–28.

178. Pocivavsek A, Burns MP, Rebeck GW. Low density lipoprotein receptors regulate microglial inflammation through c Jun N terminal kinase. Glia 2009;57(4):444–53.

179. Laskowitz DT, Wang H, Chen T, et al. Neuroprotective pentapeptide CN-105 is associated with reduced sterile inflammation and improved functional outcomes in a traumatic brain injury murine model. Scientific reports 2017;7(1):46461.

180. Vitek M, Christensen D, Wilcock D, et al. APOE-mimetic peptides reduce behavioral deficits, plaques and tangles in Alzheimer’s disease transgenics. Neurodegenerative Diseases 2012;10(1-4):122–26.

181. Contaldo C, Högger DC, Borozadi MK, et al. Radial pressure waves mediate apoptosis and functional angiogenesis during wound repair in ApoE deficient mice. Microvascular research 2012;84(1):24–33.

182. Main BS, Villapol S, Sloley SS, et al. Apolipoprotein E4 impairs spontaneous blood brain barrier repair following traumatic brain injury. Molecular neurodegeneration 2018;13:1–18.

183. Kang J, Albadawi H, Patel VI, et al. Apolipoprotein E–/–mice have delayed skeletal muscle healing after hind limb ischemia–reperfusion. Journal of vascular surgery 2008;48(3):701–08.

184. Parhizkar S, Holtzman DM. APOE mediated neuroinflammation and neurodegeneration in Alzheimer’s disease. Seminars in Immunology 2022;59:101594. doi: 10.1016/j.smim.2022.101594

185. Jordan J, Galindo MF, Miller RJ, et al. Isoform-specific effect of apolipoprotein E on cell survival and β-amyloid-induced toxicity in rat hippocampal pyramidal neuronal cultures. Journal of Neuroscience 1998;18(1):195–204.

186. Lee Y, Aono M, Laskowitz D, et al. Apolipoprotein E protects against oxidative stress in mixed neuronal–glial cell cultures by reducing glutamate toxicity. Neurochemistry international 2004;44(2):107–18.

187. Hashimoto Y, Jiang H, Niikura T, et al. Neuronal apoptosis by apolipoprotein E4 through low-density lipoprotein receptor-related protein and heterotrimeric GTPases. Journal of Neuroscience 2000;20(22):8401–09.

188. Michikawa M, Yanagisawa K. Apolipoprotein E4 induces neuronal cell death under conditions of suppressed de novo cholesterol synthesis. Journal of neuroscience research 1998;54(1):58–67.

189. Aono M, Lee Y, Grant ER, et al. Apolipoprotein E protects against NMDA excitotoxicity. Neurobiology of disease 2002;11(1):214–20.

190. Verghese PB, Castellano JM, Garai K, et al. ApoE influences amyloid-β (Aβ) clearance despite minimal apoE/Aβ association in physiological conditions. Proc Natl Acad Sci U S A 2013;110(19):E1807–16. doi: 10.1073/pnas.1220484110 [published Online First: 20130425]

191. Tai LM, Mehra S, Shete V, et al. Soluble apoE/Aβ complex: mechanism and therapeutic target for APOE4-induced AD risk. Mol Neurodegener 2014;9:2. doi: 10.1186/1750-1326-9-2 [published Online First: 20140104]

192. Kim J, Jiang H, Park S, et al. Haploinsufficiency of human APOE reduces amyloid deposition in a mouse model of amyloid-β amyloidosis. J Neurosci 2011;31(49):18007–12. doi: 10.1523/jneurosci.3773-11.2011

193. Bales KR, Liu F, Wu S, et al. Human APOE isoform-dependent effects on brain β-amyloid levels in PDAPP transgenic mice. Journal of Neuroscience 2009;29(21):6771–79.

194. Drzezga A, Grimmer T, Henriksen G, et al. Effect of APOE genotype on amyloid plaque load and gray matter volume in Alzheimer disease. Neurology 2009;72(17):1487–94.

195. Koffie RM, Hashimoto T, Tai H-C, et al. Apolipoprotein E4 effects in Alzheimer’s disease are mediated by synaptotoxic oligomeric amyloid-β. Brain 2012;135(7):2155–68.

196. Hashimoto T, Serrano-Pozo A, Hori Y, et al. Apolipoprotein E, especially apolipoprotein E4, increases the oligomerization of amyloid β peptide. Journal of Neuroscience 2012;32(43):15181–92.

197. Pillot T, Goethals M, Vanloo B, et al. Specific modulation of the fusogenic properties of the Alzheimer β amyloid peptide by apolipoprotein E isoforms. European journal of biochemistry 1997;243(3):650–59.

198. Wisniewski T, Golabek A, Matsubara E, et al. Apolipoprotein E: Binding to Soluble Alzheimer′s β-Amyloid. Biochemical and Biophysical Research Communications 1993;192(2):359–65. doi: 10.1006/bbrc.1993.1423

199. Sanan DA, Weisgraber K, Russell S, et al. Apolipoprotein E associates with beta amyloid peptide of Alzheimer’s disease to form novel monofibrils. Isoform apoE4 associates more efficiently than apoE3. The Journal of clinical investigation 1994;94(2):860–69.

200. Castaño EM, Prelli F, Pras M, et al. Apolipoprotein E Carboxyl-terminal Fragments Are Complexed to Amyloids A and L: IMPLICATIONS FOR AMYLOIDOGENESIS AND ALZHEIMER’S DISEASE *. Journal of Biological Chemistry 1995;270(29):17610–15. doi: 10.1074/jbc.270.29.17610

201. Näslund J, Thyberg J, Tjernberg LO, et al. Characterization of stable complexes involving apolipoprotein E and the amyloid beta peptide in Alzheimer’s disease brain. Neuron 1995;15(1):219–28. doi: 10.1016/0896-6273(95)90079-9

202. Chan W, Fornwald J, Brawner M, et al. Native complex formation between apolipoprotein E isoforms and the Alzheimer’s disease peptide Aβ. Biochemistry 1996;35(22):7123–30.

203. LaDu MJ, Falduto MT, Manelli AM, et al. Isoform-specific binding of apolipoprotein E to beta-amyloid. Journal of Biological Chemistry 1994;269(38):23403–06.

204. Zhou Z, Smith JD, Greengard P, et al. Alzheimer Amyloid-β Peptide Forms Denaturant-Resistant Complex with Type ε 3 but Not Type ε 4 Isoform of Native Apolipoprotein E. Molecular Medicine 1996;2(2):175–80.

205. LaDu MJ, Lukens JR, Reardon CA, et al. Association of human, rat, and rabbit apolipoprotein E with β amyloid. Journal of neuroscience research 1997;49(1):9–18.

206. Yang DS, Smith JD, Zhou Z, et al. Characterization of the binding of amyloid β peptide to cell culture derived native apolipoprotein E2, E3, and E4 isoforms and to isoforms from human plasma. Journal of neurochemistry 1997;68(2):721–25.

207. Aleshkov S, Abraham CR, Zannis VI. Interaction of Nascent ApoE2, ApoE3, and ApoE4 Isoforms Expressed in Mammalian Cells with Amyloid Peptide β (1− 40). Relevance to Alzheimer’s Disease. Biochemistry 1997;36(34):10571–80.

208. Drouet B, Fifre A, Pinçon Raymond M, et al. ApoE protects cortical neurones against neurotoxicity induced by the non fibrillar C terminal domain of the amyloid β peptide. Journal of neurochemistry 2001;76(1):117–27.

209. Nagy Z, Esiri M, Jobst K, et al. Influence of the apolipoprotein E genotype on amyloid deposition and neurofibrillary tangle formation in Alzheimer’s disease. Neuroscience 1995;69(3):757–61.

210. Sabbagh MN, Malek-Ahmadi M, Dugger BN, et al. The influence of Apolipoprotein E genotype on regional pathology in Alzheimer’s disease. BMC Neurology 2013;13(1):44. doi: 10.1186/1471-2377-13-44

211. Baek MS, Cho H, Lee HS, et al. Effect of APOE ε4 genotype on amyloid-β and tau accumulation in Alzheimer’s disease. Alzheimer’s Research & Therapy 2020;12(1):140. doi: 10.1186/s13195-020-00710-6

212. Mattsson N, Ossenkoppele R, Smith R, et al. Greater tau load and reduced cortical thickness in APOE ε4-negative Alzheimer’s disease: a cohort study. Alzheimer’s Research & Therapy 2018;10(1):77. doi: 10.1186/s13195-018-0403-x

213. Williams T, Ruiz AJ, Ruiz AM, et al. Impact of APOE genotype on prion-type propagation of tauopathy. Acta Neuropathologica Communications 2022;10(1):57. doi: 10.1186/s40478-022-01359-y

214. Chakrabarty P, Angelle C. New mechanisms highlight the complex relationship of Apolipoprotein E and tau pathogenesis. Neuron 2025;113(5):646–48. doi: 10.1016/j.neuron.2025.01.025

215. Chen G, Wang M, Zhang Z, et al. ApoE3 R136S binds to Tau and blocks its propagation, suppressing neurodegeneration in mice with Alzheimer&#x2019;s disease. Neuron doi: 10.1016/j.neuron.2024.12.015

216. Tran KM, Kwang NE, Butler CA, et al. APOE Christchurch enhances a disease-associated microglial response to plaque but suppresses response to tau pathology. Molecular Neurodegeneration 2025;20(1):9. doi: 10.1186/s13024-024-00793-x

217. Vanherle S, Janssen A, Gutiérrez de Ravé M, et al. APOE deficiency inhibits amyloid-facilitated (A) tau pathology (T) and neurodegeneration (N), halting progressive ATN pathology in a preclinical model. Molecular Psychiatry 2025 doi: 10.1038/s41380-025-03036-7

218. Blanchard JW, Bula M, Davila-Velderrain J, et al. Reconstruction of the human blood–brain barrier in vitro reveals a pathogenic mechanism of APOE4 in pericytes. Nature Medicine 2020;26(6):952–63. doi: 10.1038/s41591-020-0886-4

219. Blumenfeld J, Yip O, Kim MJ, et al. Cell type-specific roles of APOE4 in Alzheimer disease. Nature Reviews Neuroscience 2024;25(2):91–110. doi: 10.1038/s41583-023-00776-9

220. Li Z, Martens YA, Ren Y, et al. “*APOE* genotype determines cell-type-specific pathological landscape of Alzheimer’s disease. Neuron 2025;113(9):1380–97.e7. doi: 10.1016/j.neuron.2025.02.017

221. Mahley RW, Huang Y. Apolipoprotein e sets the stage: response to injury triggers neuropathology. Neuron 2012;76(5):871–85. doi: 10.1016/j.neuron.2012.11.020

222. Jiang WI, Cao Y, Xue Y, et al. Suppressing APOE4-induced neural pathologies by targeting the VHL–HIF axis. Proceedings of the National Academy of Sciences 2025;122(5):e2417515122. doi: 10.1073/pnas.2417515122

223. Wang S, Li B, Li J, et al. Cellular senescence induced by cholesterol accumulation is mediated by lysosomal ABCA1 in APOE4 and AD. Molecular Neurodegeneration 2025;20(1):15. doi: 10.1186/s13024-025-00802-7

224. Nuriel T, Peng KY, Ashok A, et al. The Endosomal–Lysosomal Pathway Is Dysregulated by APOE4 Expression in Vivo. Frontiers in Neuroscience 2017;Volume 11 - 2017 doi: 10.3389/fnins.2017.00702

225. Ellis D, Watanabe K, Wilmanski T, et al. *APOE* Genotype and Biological Age Impact Inter-Omic Associations Related to Bioenergetics. *bioRxiv* 2024:2024.10.17.618322. doi: 10.1101/2024.10.17.618322

226. Peng KY, Pérez-González R, Alldred MJ, et al. Apolipoprotein E4 genotype compromises brain exosome production. Brain 2018;142(1):163–75. doi: 10.1093/brain/awy289

227. Morrison JH, Hof PR. Life and death of neurons in the aging brain. Science 1997;278(5337):412–9. [published Online First: 1997/10/23]

228. Bussiere T, Giannakopoulos P, Bouras C, et al. Progressive degeneration of nonphosphorylated neurofilament protein-enriched pyramidal neurons predicts cognitive impairment in Alzheimer’s disease: stereologic analysis of prefrontal cortex area 9. J Comp Neurol 2003;463(3):281–302. doi: 10.1002/cne.10760 [published Online First: 2003/06/24]

229. Bussiere T, Gold G, Kovari E, et al. Stereologic analysis of neurofibrillary tangle formation in prefrontal cortex area 9 in aging and Alzheimer’s disease. Neuroscience 2003;117(3):577–92. doi: 10.1016/s0306-4522(02)00942-9 [published Online First: 2003/03/06]

230. Fu H, Possenti A, Freer R, et al. A tau homeostasis signature is linked with the cellular and regional vulnerability of excitatory neurons to tau pathology. Nat Neurosci 2019;22(1):47–56. doi: 10.1038/s41593-018-0298-7 [published Online First: 2018/12/19]

231. Ranasinghe KG, Verma P, Cai C, et al. Altered excitatory and inhibitory neuronal subpopulation parameters are distinctly associated with tau and amyloid in Alzheimer’s disease. Elife 2022;11 doi: 10.7554/eLife.77850 [published Online First: 20220526]

232. Chen S, Chang Y, Li L, et al. Spatially resolved transcriptomics reveals genes associated with the vulnerability of middle temporal gyrus in Alzheimer’s disease. Acta Neuropathol Commun 2022;10(1):188. doi: 10.1186/s40478-022-01494-6 [published Online First: 20221221]

233. Goedert M, Wischik C, Crowther R, et al. Cloning and sequencing of the cDNA encoding a core protein of the paired helical filament of Alzheimer disease: identification as the microtubule-associated protein tau. Proceedings of the National Academy of Sciences 1988;85(11):4051–55.

234. Spires-Jones TL, Stoothoff WH, De Calignon A, et al. Tau pathophysiology in neurodegeneration: a tangled issue. Trends in neurosciences 2009;32(3):150–59.

235. Spires-Jones TL, Kopeikina KJ, Koffie RM, et al. Are tangles as toxic as they look? J Mol Neurosci 2011;45(3):438–44. doi: 10.1007/s12031-011-9566-7 [published Online First: 20110603]

236. Delacourte A, Buée L. Normal and pathological Tau proteins as factors for microtubule assembly. Int Rev Cytol 1997;171:167–224. doi: 10.1016/s0074-7696(08)62588-7

237. Brion J-P. Neurofibrillary tangles and Alzheimer’s disease. European neurology 1998;40(3):130–40.

238. Ehrenberg A, Nguy A, Theofilas P, et al. Quantifying the accretion of hyperphosphorylated tau in the locus coeruleus and dorsal raphe nucleus: the pathological building blocks of early Alzheimer’s disease. Neuropathology and applied neurobiology 2017;43(5):393–408.

239. Grinberg L, Rüb U, Ferretti R, et al. The dorsal raphe nucleus shows phospho tau neurofibrillary changes before the transentorhinal region in Alzheimer’s disease. A precocious onset? Neuropathology and applied neurobiology 2009;35(4):406–16.

240. Grudzien A, Shaw P, Weintraub S, et al. Locus coeruleus neurofibrillary degeneration in aging, mild cognitive impairment and early Alzheimer’s disease. Neurobiology of aging 2007;28(3):327–35.

241. Tomlinson B, Irving D, Blessed G. Cell loss in the locus coeruleus in senile dementia of Alzheimer type. Journal of the neurological sciences 1981;49(3):419–28.

242. Theofilas P, Ehrenberg AJ, Dunlop S, et al. Locus coeruleus volume and cell population changes during Alzheimer’s disease progression: a stereological study in human postmortem brains with potential implication for early-stage biomarker discovery. Alzheimer’s & Dementia 2017;13(3):236–46.

243. La Joie R, Visani AV, Baker SL, et al. Prospective longitudinal atrophy in Alzheimer’s disease correlates with the intensity and topography of baseline tau-PET. Sci Transl Med 2020;12(524) doi: 10.1126/scitranslmed.aau5732 [published Online First: 2020/01/03]

244. Hanseeuw BJ, Betensky RA, Jacobs HIL, et al. Association of Amyloid and Tau With Cognition in Preclinical Alzheimer Disease: A Longitudinal Study. JAMA Neurol 2019;76(8):915–24. doi: 10.1001/jamaneurol.2019.1424 [published Online First: 2019/06/04]

245. Haroutunian V, Purohit DP, Perl DP, et al. Neurofibrillary tangles in nondemented elderly subjects and mild Alzheimer disease. Archives of neurology 1999;56(6):713–18.

246. Braak H, Braak E. Neuropathological stageing of Alzheimer-related changes. Acta neuropathologica 1991;82(4):239–59.

247. Bierer LM, Hof PR, Purohit DP, et al. Neocortical neurofibrillary tangles correlate with dementia severity in Alzheimer’s disease. Archives of neurology 1995;52(1):81–88.

248. Duyckaerts C, Bennecib M, Grignon Y, et al. Modeling the Relation Between Neurofibrillary Tangles and Intellectual Status. Neurobiology of Aging 1997;18(3):267–73. doi: 10.1016/S0197-4580(97)80306-5

249. Arriagada PV, Growdon JH, Hedley-Whyte ET, et al. Neurofibrillary tangles but not senile plaques parallel duration and severity of Alzheimer’s disease. Neurology 1992;42(3 Pt 1):631–9. [published Online First: 1992/03/01]

250. Terry RD. Do neuronal inclusions kill the cell? Advances in Dementia Research 2000:91–93.

251. Lennol MP, Bordier C, Kamelher L, et al. ApoE-calypse tau: ApoE–tau synergy in Alzheimer’s disease. Journal of Experimental Medicine 2025;222(10) doi: 10.1084/jem.20250965

252. Bell WR, An Y, Kageyama Y, et al. Neuropathologic, genetic, and longitudinal cognitive profiles in primary age-related tauopathy (PART) and Alzheimer’s disease. Alzheimer’s & Dementia 2019;15(1):8–16. doi: 10.1016/j.jalz.2018.07.215

253. Andorfer C, Acker CM, Kress Y, et al. Cell-cycle reentry and cell death in transgenic mice expressing nonmutant human tau isoforms. Journal of Neuroscience 2005;25(22):5446–54.

254. Dunckley T, Beach TG, Ramsey KE, et al. Gene expression correlates of neurofibrillary tangles in Alzheimer’s disease. Neurobiology of Aging 2006;27(10):1359–71. doi: 10.1016/j.neurobiolaging.2005.08.013

255. Kuchibhotla KV, Wegmann S, Kopeikina KJ, et al. Neurofibrillary tangle-bearing neurons are functionally integrated in cortical circuits in vivo. Proc Natl Acad Sci U S A 2014;111(1):510–4. doi: 10.1073/pnas.1318807111 [published Online First: 20131224]

256. Rudinskiy N, Hawkes JM, Wegmann S, et al. Tau pathology does not affect experience-driven single-neuron and network-wide Arc/Arg3.1 responses. Acta Neuropathologica Communications 2014;2(1):63. doi: 10.1186/2051-5960-2-63

257. Rocher AB, Crimins JL, Amatrudo JM, et al. Structural and functional changes in tau mutant mice neurons are not linked to the presence of NFTs. Experimental Neurology 2010;223(2):385–93. doi: 10.1016/j.expneurol.2009.07.029

258. Spires-Jones TL, De Calignon A, Matsui T, et al. In vivo imaging reveals dissociation between caspase activation and acute neuronal death in tangle-bearing neurons. Journal of Neuroscience 2008;28(4):862–67.

259. de Calignon A, Spires-Jones TL, Pitstick R, et al. Tangle-Bearing Neurons Survive Despite Disruption of Membrane Integrity in a Mouse Model of Tauopathy. Journal of Neuropathology & Experimental Neurology 2009;68(7):757–61. doi: 10.1097/NEN.0b013e3181a9fc66

260. Zwang TJ, Sastre Ed, Wolf N, et al. Neurofibrillary tangle-bearing neurons have reduced risk of cell death in mice with Alzheimer&#x2019;s pathology. Cell Reports 2024;43(8) doi: 10.1016/j.celrep.2024.114574

261. Kopeikina KJ, Hyman BT, Spires-Jones TL. Soluble forms of tau are toxic in Alzheimer’s disease. Transl Neurosci 2012;3(3):223–33. doi: 10.2478/s13380-012-0032-y

262. Clavaguera F, Bolmont T, Crowther RA, et al. Transmission and spreading of tauopathy in transgenic mouse brain. Nature cell biology 2009;11(7):909–13.

263. de Calignon A, Polydoro M, Suarez-Calvet M, et al. Propagation of tau pathology in a model of early Alzheimer’s disease. Neuron 2012;73(4):685–97. doi: 10.1016/j.neuron.2011.11.033

264. Bolós M, Pallas-Bazarra N, Terreros-Roncal J, et al. Soluble Tau has devastating effects on the structural plasticity of hippocampal granule neurons. Transl Psychiatry 2017;7(12):1267. doi: 10.1038/s41398-017-0013-6 [published Online First: 20171208]

265. Lasagna-Reeves CA, Castillo-Carranza DL, Sengupta U, et al. Tau oligomers impair memory and induce synaptic and mitochondrial dysfunction in wild-type mice. Molecular Neurodegeneration 2011;6(1):39. doi: 10.1186/1750-1326-6-39

266. Sanchez-Mejias E, Navarro V, Jimenez S, et al. Soluble phospho-tau from Alzheimer’s disease hippocampus drives microglial degeneration. Acta Neuropathologica 2016;132(6):897–916. doi: 10.1007/s00401-016-1630-5

267. Richardson TE, Orr ME, Orr TC, et al. Spatial proteomic differences in chronic traumatic encephalopathy, Alzheimer’s disease, and primary age related tauopathy hippocampi. Alzheimer’s & Dementia 2025;21(2):e14487.

268. Jurga AM, Paleczna M, Kadluczka J, et al. Beyond the GFAP-Astrocyte Protein Markers in the Brain. Biomolecules 2021;11(9) doi: 10.3390/biom11091361 [published Online First: 20210914]

269. Hol EM, Roelofs RF, Moraal E, et al. Neuronal expression of GFAP in patients with Alzheimer pathology and identification of novel GFAP splice forms. Molecular Psychiatry 2003;8(9):786–96. doi: 10.1038/sj.mp.4001379

270. Doetsch F, Caille I, Lim DA, et al. Subventricular zone astrocytes are neural stem cells in the adult mammalian brain. Cell 1999;97(6):703–16.

271. Laywell ED, Rakic P, Kukekov VG, et al. Identification of a multipotent astrocytic stem cell in the immature and adult mouse brain. Proceedings of the National Academy of Sciences 2000;97(25):13883–88.

272. Seri B, Garcıa-Verdugo JM, McEwen BS, et al. Astrocytes give rise to new neurons in the adult mammalian hippocampus. Journal of Neuroscience 2001;21(18):7153–60.

273. Rahman MW, Sharma P, Chattopadhyay T, et al. Distinct neuronal vulnerability and metabolic dysfunctions are characteristic features of fast-progressing Alzheimer’s patients with Lewy bodies. Journal of Biological Chemistry 2025;301(4) doi: 10.1016/j.jbc.2025.108396

274. Calvo-Rodriguez M, Hou SS, Snyder AC, et al. Increased mitochondrial calcium levels associated with neuronal death in a mouse model of Alzheimer’s disease. Nature Communications 2020;11(1):2146. doi: 10.1038/s41467-020-16074-2

275. Mrdjen D, Fox EJ, Bukhari SA, et al. The basis of cellular and regional vulnerability in Alzheimer’s disease. Acta Neuropathol 2019;138(5):729–49. doi: 10.1007/s00401-019-02054-4 [published Online First: 20190807]

276. Hof PR, Nimchinsky EA, Celio MR, et al. Calretinin-immunoreactive neocortical interneurons are unaffected in Alzheimer’s disease. Neuroscience letters 1993;152(1-2):145–49.

277. Sampson VL, Morrison JH, Vickers JC. The cellular basis for the relative resistance of parvalbumin and calretinin immunoreactive neocortical neurons to the pathology of Alzheimer’s disease. Experimental neurology 1997;145(1):295–302.

278. Mattson MP, Cheng B, Davis D, et al. beta-Amyloid peptides destabilize calcium homeostasis and render human cortical neurons vulnerable to excitotoxicity. Journal of Neuroscience 1992;12(2):376–89.

279. Mrdjen D, Cannon BJ, Amouzgar M, et al. Spatial proteomics of Alzheimer’s disease-specific human microglial states. Nature Immunology 2025;26(8):1397–410. doi: 10.1038/s41590-025-02203-w

280. Thrupp N, Sala Frigerio C, Wolfs L, et al. Single-Nucleus RNA-Seq Is Not Suitable for Detection of Microglial Activation Genes in Humans. Cell Rep 2020;32(13):108189. doi: 10.1016/j.celrep.2020.108189

281. Podlesny-Drabiniok A, Novikova G, Liu Y, et al. BHLHE40/41 regulate macrophage/microglia responses associated with Alzheimer’s disease and other disorders of lipid-rich tissues. bioRxiv 2023:2023.02.13.528372. doi: 10.1101/2023.02.13.528372

282. Ardura-Fabregat A, Bosch LFP, Wogram E, et al. Response of spatially defined microglia states with distinct chromatin accessibility in a mouse model of Alzheimer’s disease. Nature Neuroscience 2025 doi: 10.1038/s41593-025-02006-0

283. Tieu T, Cruz A-JN, Weinstein JR, et al. Physiological and injury-induced microglial dynamics across the lifespan. Cell Reports 2025;44(7) doi: 10.1016/j.celrep.2025.115991

284. Guvenek A, Parikshak N, Zamolodchikov D, et al. Transcriptional profiling in microglia across physiological and pathological states identifies a transcriptional module associated with neurodegeneration. Communications Biology 2024;7(1):1168. doi: 10.1038/s42003-024-06684-7

285. Perea JR, Bolós M, Avila J. Microglia in Alzheimer’s Disease in the Context of Tau Pathology. Biomolecules 2020;10(10) doi: 10.3390/biom10101439 [published Online First: 20201014]

286. Zhang Y, Chen K, Sloan SA, et al. An RNA-sequencing transcriptome and splicing database of glia, neurons, and vascular cells of the cerebral cortex. Journal of neuroscience 2014;34(36):11929–47.

287. Zhang Y, Sloan SA, Clarke LE, et al. Purification and characterization of progenitor and mature human astrocytes reveals transcriptional and functional differences with mouse. Neuron 2016;89(1):37–53.

288. Wendeln A-C, Degenhardt K, Kaurani L, et al. Innate immune memory in the brain shapes neurological disease hallmarks. Nature 2018;556(7701):332–38.

289. March-Diaz R, Lara-Ureña N, Romero-Molina C, et al. Hypoxia compromises the mitochondrial metabolism of Alzheimer’s disease microglia via HIF1. Nature Aging 2021;1(4):385–99. doi: 10.1038/s43587-021-00054-2

290. Lu D-Y, Liou H-C, Tang C-H, et al. Hypoxia-induced iNOS expression in microglia is regulated by the PI3-kinase/Akt/mTOR signaling pathway and activation of hypoxia inducible factor-1α. Biochemical Pharmacology 2006;72(8):992–1000. doi: 10.1016/j.bcp.2006.06.038

291. Holmes BB, Weigel TK, Chung JM, et al. β-Amyloid Induces Microglial Expression of GPC4 and APOE Leading to Increased Neuronal Tau Pathology and Toxicity. bioRxiv 2025:2025.02.20.637701. doi: 10.1101/2025.02.20.637701

292. van Olst L, Simonton B, Edwards AJ, et al. Microglial mechanisms drive amyloid-β clearance in immunized patients with Alzheimer’s disease. Nature Medicine 2025;31(5):1604–16. doi: 10.1038/s41591-025-03574-1

293. Dolan M-J, Therrien M, Jereb S, et al. Exposure of iPSC-derived human microglia to brain substrates enables the generation and manipulation of diverse transcriptional states in vitro. Nature Immunology 2023;24(8):1382–90. doi: 10.1038/s41590-023-01558-2

294. Kosoy R, Fullard JF, Bendl J, et al. Alzheimer’s disease transcriptional landscape in ex vivo human microglia. Nature Neuroscience 2025 doi: 10.1038/s41593-025-02020-2

295. Gao C, Jiang J, Tan Y, et al. Microglia in neurodegenerative diseases: mechanism and potential therapeutic targets. Signal Transduction and Targeted Therapy 2023;8(1):359. doi: 10.1038/s41392-023-01588-0

296. Xie M, Liu YU, Zhao S, et al. TREM2 interacts with TDP-43 and mediates microglial neuroprotection against TDP-43-related neurodegeneration. Nature neuroscience 2022;25(1):26–38.

297. Bemiller SM, McCray TJ, Allan K, et al. TREM2 deficiency exacerbates tau pathology through dysregulated kinase signaling in a mouse model of tauopathy. Molecular neurodegeneration 2017;12:1–12.

298. Jay TR, Hirsch AM, Broihier ML, et al. Disease Progression-Dependent Effects of TREM2 Deficiency in a Mouse Model of Alzheimer’s Disease. The Journal of Neuroscience 2017;37(3):637–47. doi: 10.1523/jneurosci.2110-16.2016

299. Rachmian N, Medina S, Cherqui U, et al. Identification of senescent, TREM2-expressing microglia in aging and Alzheimer’s disease model mouse brain. Nature Neuroscience 2024;27(6):1116–24. doi: 10.1038/s41593-024-01620-8

300. Ikenouchi J, Umeda M. FRMD4A regulates epithelial polarity by connecting Arf6 activation with the PAR complex. Proceedings of the National Academy of Sciences 2010;107(2):748–53. doi: 10.1073/pnas.0908423107

301. Tepass U. FERM proteins in animal morphogenesis. Current Opinion in Genetics & Development 2009;19(4):357–67. doi: 10.1016/j.gde.2009.05.006

302. Lambert JC, Grenier-Boley B, Harold D, et al. Genome-wide haplotype association study identifies the FRMD4A gene as a risk locus for Alzheimer’s disease. Mol Psychiatry 2013;18(4):461–70. doi: 10.1038/mp.2012.14 [published Online First: 2012/03/21]

303. Andrews JS, Desai U, Kirson NY, et al. Disease severity and minimal clinically important differences in clinical outcome assessments for Alzheimer’s disease clinical trials. Alzheimers Dement (N Y) 2019;5:354–63. doi: 10.1016/j.trci.2019.06.005 [published Online First: 20190802]

304. Uronen R-LE, Huttunen HJ. Genetic risk factors of Alzheimer’s disease and cell-to-cell transmission of Tau. The Journal of Neurology and Neuromedicine 2016 doi: 10.29245/2572.942X/2016/2.1022

305. Martiskainen H, Viswanathan J, Nykänen N-P, et al. Transcriptomics and mechanistic elucidation of Alzheimer’s disease risk genes in the brain and in vitro models. Neurobiology of Aging 2015;36(2):1221.e15-21.e28. doi: 10.1016/j.neurobiolaging.2014.09.003

306. Yan X, Nykänen NP, Brunello CA, et al. FRMD4A-cytohesin signaling modulates the cellular release of tau. J Cell Sci 2016;129(10):2003–15. doi: 10.1242/jcs.180745 [published Online First: 20160404]

307. Insel PS, Hansson O, Mattsson-Carlgren N. Association Between Apolipoprotein E ε2 vs ε4, Age, and β-Amyloid in Adults Without Cognitive Impairment. JAMA Neurology 2021;78(2):229–35. doi: 10.1001/jamaneurol.2020.3780

308. Hanamsagar R, Alter MD, Block CS, et al. Generation of a microglial developmental index in mice and in humans reveals a sex difference in maturation and immune reactivity. Glia 2017;65(9):1504–20.

309. Lake BB, Codeluppi S, Yung YC, et al. A comparative strategy for single-nucleus and single-cell transcriptomes confirms accuracy in predicted cell-type expression from nuclear RNA. Scientific reports 2017;7(1):6031.

310. Yu G, Thorpe A, Zeng Q, et al. The Landscape of Sex- and APOE Genotype-Specific Transcriptional Changes in Alzheimer’s Disease at the Single Cell Level. bioRxiv 2024:2024.12.01.626234. doi: 10.1101/2024.12.01.626234

311. Harerimana NV, Goate AM, Bowles KR. The influence of 17q21.31 and APOE genetic ancestry on neurodegenerative disease risk. Frontiers in Aging Neuroscience 2022;Volume 14 - 2022 doi: 10.3389/fnagi.2022.1021918

312. Beydoun MA, Weiss J, Beydoun HA, et al. Race, APOE genotypes, and cognitive decline among middle-aged urban adults. Alzheimer’s Research & Therapy 2021;13(1):120.

313. Beydoun MA, Boueiz A, Abougergi MS, et al. Sex differences in the association of the apolipoprotein E epsilon 4 allele with incidence of dementia, cognitive impairment, and decline. Neurobiology of Aging 2012;33(4):720–31.e4. doi: 10.1016/j.neurobiolaging.2010.05.017

314. Bakken TE, Hodge RD, Miller JA, et al. Single-nucleus and single-cell transcriptomes compared in matched cortical cell types. PloS one 2018;13(12):e0209648.

315. Ding J, Adiconis X, Simmons SK, et al. Systematic comparison of single-cell and single-nucleus RNA-sequencing methods. Nature Biotechnology 2020;38(6):737–46. doi: 10.1038/s41587-020-0465-8

316. Lacar B, Linker SB, Jaeger BN, et al. Nuclear RNA-seq of single neurons reveals molecular signatures of activation. Nature communications 2016;7(1):11022.

317. Wang Y, Fan JL, Melms JC, et al. Multimodal single-cell and whole-genome sequencing of small, frozen clinical specimens. Nature genetics 2023;55(1):19–25.

318. van den Brink SC, Sage F, Vértesy Á, et al. Single-cell sequencing reveals dissociation-induced gene expression in tissue subpopulations. Nature methods 2017;14(10):935–36.

319. Jiang A, Lehnert K, Reid SJ, et al. Isolated nuclei from frozen tissue are the superior source for single cell RNA-seq compared with whole cells. bioRxiv 2023:2023.02. 19.529150.

320. Denisenko E, Guo BB, Jones M, et al. Systematic assessment of tissue dissociation and storage biases in single-cell and single-nucleus RNA-seq workflows. Genome biology 2020;21:1–25.

321. Kuo A, Hansen KD, Hicks SC. Quantification and statistical modeling of droplet-based single-nucleus RNA-sequencing data. Biostatistics 2024;25(3):801–17.

322. Chamberlin JT, Lee Y, Marth GT, et al. Differences in molecular sampling and data processing explain variation among single-cell and single-nucleus RNA-seq experiments. Genome Res 2024;34(2):179–88. doi: 10.1101/gr.278253.123 [published Online First: 20240320]

323. Pettas S, Karagianni K, Kanata E, et al. Profiling Microglia through Single-Cell RNA Sequencing over the Course of Development, Aging, and Disease. Cells 2022;11(15) doi: 10.3390/cells11152383 [published Online First: 20220802]

324. Cosentino S, Scarmeas N, Helzner E, et al. APOE epsilon 4 allele predicts faster cognitive decline in mild Alzheimer disease. Neurology 2008;70(19 Pt 2):1842-9. doi: 10.1212/01.wnl.0000304038.37421.cc [published Online First: 20080409]

325. Martins CA, Oulhaj A, de Jager CA, et al. APOE alleles predict the rate of cognitive decline in Alzheimer disease: a nonlinear model. Neurology 2005;65(12):1888–93. doi: 10.1212/01.wnl.0000188871.74093.12

326. Craft S, Teri L, Edland SD, et al. Accelerated decline in apolipoprotein E-epsilon4 homozygotes with Alzheimer’s disease. Neurology 1998;51(1):149–53. doi: 10.1212/wnl.51.1.149

327. Hirono N, Hashimoto M, Yasuda M, et al. Accelerated memory decline in Alzheimer’s disease with apolipoprotein epsilon4 allele. J Neuropsychiatry Clin Neurosci 2003;15(3):354–8. doi: 10.1176/jnp.15.3.354

328. Kanai M, Shizuka M, Urakami K, et al. Apolipoprotein E4 accelerates dementia and increases cerebrospinal fluid tau levels in Alzheimer’s disease. Neurosci Lett 1999;267(1):65–8. doi: 10.1016/s0304-3940(99)00323-7

329. Chang YL, Fennema-Notestine C, Holland D, et al. APOE interacts with age to modify rate of decline in cognitive and brain changes in Alzheimer’s disease. Alzheimers Dement 2014;10(3):336–48. doi: 10.1016/j.jalz.2013.05.1763 [published Online First: 20130727]

330. Morrison C, Oliver MD, Berry V, et al. The influence of APOE status on rate of cognitive decline. GeroScience 2024:1–12.

331. Qian J, Zhang Y, Betensky RA, et al. Neuropathology-Independent Association Between *APOE* Genotype and Cognitive Decline Rate in the Normal Aging-Early Alzheimer Continuum. Neurology Genetics 2023;9(1):e200055. doi: doi:10.1212/NXG.0000000000200055

332. Stern Y, Brandt J, Albert M, et al. The absence of an apolipoprotein epsilon4 allele is associated with a more aggressive form of Alzheimer’s disease. Ann Neurol 1997;41(5):615–20. doi: 10.1002/ana.410410510

333. Frisoni GB, Govoni S, Geroldi C, et al. Gene dose of the epsilon 4 allele of apolipoprotein E and disease progression in sporadic late-onset Alzheimer’s disease. Ann Neurol 1995;37(5):596–604. doi: 10.1002/ana.410370509

334. Hoyt BD, Massman PJ, Schatschneider C, et al. Individual growth curve analysis of APOE epsilon 4-associated cognitive decline in Alzheimer disease. Arch Neurol 2005;62(3):454–9. doi: 10.1001/archneur.62.3.454

335. Kleiman T, Zdanys K, Black B, et al. Apolipoprotein E epsilon4 allele is unrelated to cognitive or functional decline in Alzheimer’s disease: retrospective and prospective analysis. Dement Geriatr Cogn Disord 2006;22(1):73–82. doi: 10.1159/000093316 [published Online First: 20060512]

336. Growdon JH, Locascio JJ, Corkin S, et al. Apolipoprotein E genotype does not influence rates of cognitive decline in Alzheimer’s disease. Neurology 1996;47(2):444–8. doi: 10.1212/wnl.47.2.444

337. Holmes C, Levy R, McLoughlin DM, et al. Apolipoprotein E: non-cognitive symptoms and cognitive decline in late onset Alzheimer’s disease. J Neurol Neurosurg Psychiatry 1996;61(6):580–3. doi: 10.1136/jnnp.61.6.580

338. Kurz A, Egensperger R, Haupt M, et al. Apolipoprotein E epsilon 4 allele, cognitive decline, and deterioration of everyday performance in Alzheimer’s disease. Neurology 1996;47(2):440–3. doi: 10.1212/wnl.47.2.440

339. Basun H, Grut M, Winblad B, et al. Apolipoprotein epsilon 4 allele and disease progression in patients with late-onset Alzheimer’s disease. Neurosci Lett 1995;183(1-2):32–4. doi: 10.1016/0304-3940(94)11107-t

340. Farlow MR, Cyrus PA, Nadel A, et al. Metrifonate treatment of AD: influence of APOE genotype. Neurology 1999;53(9):2010–6. doi: 10.1212/wnl.53.9.2010

341. Aerssens J, Raeymaekers P, Lilienfeld S, et al. APOE genotype: no influence on galantamine treatment efficacy nor on rate of decline in Alzheimer’s disease. Dement Geriatr Cogn Disord 2001;12(2):69–77. doi: 10.1159/000051238

342. Murphy GM, Jr., Taylor J, Kraemer HC, et al. No association between apolipoprotein E epsilon 4 allele and rate of decline in Alzheimer’s disease. Am J Psychiatry 1997;154(5):603–8. doi: 10.1176/ajp.154.5.603

343. Polsinelli AJ, Lane KA, Manchella MK, et al. APOE ε4 is associated with earlier symptom onset in LOAD but later symptom onset in EOAD. Alzheimers Dement 2023;19(5):2212–17. doi: 10.1002/alz.12955 [published Online First: 20230201]

